# Circadian disruption alters hepatic calcium hemostasis, endocannabinoidome and mitochondria through *N*-docosahexaenoyl ethanolamide-GPR110 signaling

**DOI:** 10.64898/2026.06.10.731470

**Authors:** Pejman A. Pashaki, Timothy Niepokny, Elizabeth Dumais, Eric M. Mintz, David Marsolais, Alain Veilleux, Nicolas Flamand, Vincenzo Di Marzo, Cristoforo Silvestri

## Abstract

Circadian rhythm disruption is associated with metabolic and inflammatory disorders; however, the mechanisms linking circadian dysfunction to endocannabinoidome (eCBome) signaling and mitochondrial metabolism remain unclear. In our previous *in vivo* study, constant light exposure altered hepatic eCBome profiles, reduced *N*-acylethanolamines (NAEs), increased monoacylglycerols (MAGs), and elevated inflammatory cytokines. Here, we investigated the underlying mechanisms using CRISPR/Cas9-generated BMAL1 knockout (KO) HepG2 cells as an *in vitro* model of circadian alteration. The BMAL1 KO model showed broad lipid remodeling characterized by increased fatty acids, prostaglandins, and MAGs together with reduced NAEs and enhanced lipid accumulation. These changes were accompanied by increased inflammatory signaling and cytokine production. Among the assessed genes, *GPR110* was significantly altered in mice exposed to constant light (*in vivo* study) and BMAL1 KO model and emerged as a potential mediator linking circadian signaling to mitochondrial function. BMAL1 KO cells also exhibited significantly increased calcium (Ca²^+^) levels in mitochondria and the endoplasmic reticulum (ER), along with attenuation of mitochondrial and glycolytic ATP production. BMAL1KO did not abolish the rhythmicity of NAEs level over 24 hours from medium deprivation and read ministration except for *N*-docosahexaenoyl-ethanolamide (DHEA). Further, experiments showed that DHEA acts through GPR110 and suppress inflammatory lipid-associated pathways, enhances ATP production, and increases mitochondrial and ER Ca²^+^ accumulation and inflammatory signaling. Together, these mitochondrial Ca²^+^ signaling, and inflammation in hepatocytes, highlighting DHEA-GPR110 signaling as a potential regulator of hepatic metabolic homeostasis.

**Highlights:** Circadian disruption increases hepatic monoacylglycerols and decreases *N*-acylethanolamines.

Circadian disruption decreases ATP production and enhances mitochondrial and endoplasmic reticulum Ca²^+^ levels in hepatocytes

DHEA-GPR110 signaling regulates hepatocytes mitochondrial Ca²^+^ dynamics and ATP production

GPR110-mediated Ca²^+^ signaling significantly alters hepatocytes glycolysis and glycolytic ATP production

## Introduction

Circadian clocks are an endogenous timekeeping system that synchronizes metabolic processes with environmental cues (1). In liver, this system coordinates oscillations in genes involved in lipid metabolism, mitochondrial activity, and redox balance, thereby aligning metabolic homeostasis with feeding-fasting and energy cycles (2). Core clock genes include brain and muscle ARNT-like protein 1 (BMAL1), circadian locomotor output cycles kaput (Clock), Period (PER1, PER2, PER3), Cryptochrome (CRY1, CRY2), Retinoic acid receptor-related orphan receptors (ROR), and REV-ERBα (also known as NR1D1; nuclear receptor subfamily 1 group D member 1). These genes encode for protein that together form interconnected transcriptional-translational feedback loops in which the CLOCK:BMAL1 heterodimer binds E-box enhancer elements to activate transcription of target genes including PER, CRY, ROR, and REV-ERB, while accumulated PER and CRY proteins subsequently inhibit CLOCK:BMAL1 activity to repress their own expression. ROR and REV-ERB oppositely regulate BMAL1 transcription through activation and repression, respectively, thereby generating rhythmic oscillations that sustain circadian rhythmicity (3).

Circadian disruption increases body weight (4) as well hepatic lipid accumulation and is associated with inflammation (5). This inflammatory state is typically characterized as chronic low-grade inflammation, which can exacerbate inflammatory responses through dysregulation of clock-controlled pathways. One mechanism underlying this phenomenon is the anti-inflammatory role of BMAL1 that suppresses pro-inflammatory signaling. Consistent with this role, reduced BMAL1 expression has been associated with increased production of inflammatory mediators such as IL-6 (6, 7). Circadian disruption may also impair mitochondrial function through two interconnected mechanisms. First, disruption of the molecular clock itself, including BMAL1 or CLOCK deficiency, reduces mitochondrial membrane potential, oxygen consumption, electron transport chain activity, and ATP production (8–11). Second, the chronic inflammatory environment induced by circadian disruption can further compromise mitochondrial bioenergetics and contribute to metabolic dysfunction (12, 13). Together, these effects suggest that impaired BMAL1 activity may promote inflammation, mitochondrial dysfunction, and lipid accumulation, thereby contributing to the metabolic disturbances associated with circadian disruption.

Mechanistically, circadian clock components including Period (PER1 and PER2) and BMAL1 contribute substantially to diurnal nutrient utilization and metabolic homeostasis, with PER1/PER2 regulating rate-limiting enzymes involved in mitochondrial metabolism and BMAL1 playing central roles in hepatic lipid metabolism (14), fatty acid oxidation, and mitochondrial dynamics (11). BMAL1 deficiency has been associated with altered fatty acid metabolism, impaired β-oxidation, mitochondrial dysfunction, oxidative stress, glucose intolerance, inflammation, and dysregulated lipogenesis (15–17). Consistent with these metabolic alterations, circadian disruption caused by either environmental stressors or genetic manipulation impairs liver function and increases susceptibility to hepatic steatosis and metabolic liver diseases (18, 19). Both global and liver-specific BMAL1 deficiency alter lipogenesis-related gene expression and promote ectopic lipid accumulation in the liver and skeletal muscle through insulin-mTORC2-AKT signaling (16, 20). However, global BMAL1 deficiency, rather than hepatocyte-specific BMAL1 deletion alone, plays a central role in regulating liver inflammation, fibrosis, and metabolic adaptation due to systemic endocrine perturbations (21). BMAL1 directly up-regulates the expression of Peroxisome Proliferator-Activated Receptor Alpha (PPARα) (22), promoting its rhythmic expression and contributing to its protective effects against alcoholic liver disease (ALD) (15). However, the relationship between BMAL1 and PPARα is bidirectional and regulate each other (23). In addition to its role in lipid metabolism, BMAL1 transcriptionally regulates mitochondrial biogenesis and mitochondrial dynamics, particularly mitochondrial fission and mitophagy, in accordance with circadian metabolic demands. Loss of BMAL1 function results in swollen and metabolically impaired mitochondria that fail to adapt to nutrient fluctuations, leading to reduced mitochondrial respiration and increased oxidative stress (11). Liver-specific BMAL1 knockout (LBMAL1KO) mice accumulate oxidative damage, whereas restoration of mitochondrial fission through FIS1 expression partially rescues mitochondrial morphological and metabolic defects (11). Another potential mechanism involved in liver metabolism and circadian rhythm is patatin-like phospholipase domain-containing protein 3 (PNPLA3), a critical regulator of lipid metabolism in hepatocytes which has rhythm and directly influences lipid droplet dynamics (24).

Recent evidence suggests that lipid mediators of the endocannabinoidome (eCBome), particularly monoacylglycerols (MAGs) and *N-*acylethanolamines (NAEs), are under circadian regulation and contribute to metabolic homeostasis (25). The canonical biosynthesis of NAEs involves two sequential enzymatic steps. First, Ca2+-dependent *N*-acyltransferase (NAT) transfers an acyl group from membrane phospholipids to phosphatidylethanolamine (PE), generating *N*-acylphosphatidylethanolamine (NAPE), a rate-limiting step strongly influenced by intracellular Ca2+ levels. Subsequently, NAPE-specific phospholipase D (NAPE-PLD) hydrolyzes NAPE to produce NAEs and phosphatidic acid, whereas NAE degradation is primarily mediated by fatty acid amide hydrolase 1 (FAAH) (26, 27). Interestingly, NAPE-PLD expression exhibits circadian rhythmicity and may be under core clock regulation (22). Similarly, MAG biosynthesis involves the sequential hydrolysis of arachidonic acid-containing phospholipids by phospholipase C (PLC) and diacylglycerol lipase (DAGL). PLC hydrolyzes phosphatidylinositol-4,5-bisphosphate to generate diacylglycerol, which is subsequently converted into MAGs by DAGL α and β isozymes that, like NAT, are influenced by Ca2+ (28, 29). MAG degradation is mainly mediated by monoacylglycerol lipase (MAGL), whose expression also displays circadian rhythmicity and may be regulated by core clock machinery (22).

In our previous study, 15 days of constant light exposure significantly altered tissue-specific eCBome profiles, particularly in the liver, where MAG levels 2-Stearidonoyl-Glycerol (2-SDG) and 2-Oleoyl-Glycerol (2-OG)) were increased, whereas the NAEs *N*-docosahexaenoyl ethanolamine (DHEA) and *N*-Linoleoyl-Ethanolamine (LEA) were reduced (30). These findings suggested a strong relationship between circadian rhythm disruption and hepatic eCBome remodeling; however, the mechanistic consequences of the alterations of these bioactive lipid mediators at the cellular level remained unknown. This question is particularly important because elevated MAG levels of endocannabinoids, anandamide and 2-Arachidonoyl-Glycerol (2-AG), and some of their respective NAE and MAG congeners, are associated with metabolic liver disorders (31), whereas several NAEs possess anti-inflammatory and hepatoprotective properties (32–35). Therefore, in the present study, we focused on the potential role of NAEs, particularly DHEA, in hepatic metabolism under circadian disruption conditions.

Among the NAEs, DHEA functions as a bioactive signaling molecule through activation of the orphan adhesion G protein-coupled receptor GPR110 (36). DHEA binds within the hydrophobic pocket of GPR110, normally occupied by the Stachel peptide in the inactive receptor state, leading to receptor activation and stimulation of cAMP/PKA/CREB signaling pathways (37–39). In addition to its signaling properties, GPR110 has been implicated in hepatic lipid metabolism and inflammatory regulation. Hepatic GPR110 expression positively correlates with liver fat accumulation in obese mice and patients with non-alcoholic fatty liver disease (NAFLD), partly through modulation of Scd1 expression (40). Furthermore, increased GPR110 expression has been associated with activation of inflammatory pathways, including IL-6/STAT3 signaling, whereas DHEA-GPR110 signaling has been linked to anti-inflammatory effects (41, 42). Together, these findings suggest that DHEA-GPR110 signaling may represent a molecular interface linking circadian disruption, lipid mediator remodeling, inflammation, and mitochondrial metabolism.

To investigate the cell-autonomous role of the circadian clock in hepatocyte metabolism, we generated BMAL1-knockout HepG2 cells as an *in vitro* model of circadian disruption. Using LC-MS/MS lipidomics, TaqMan gene expression profiling, calcium (Ca²^+^) biosensors, and Seahorse mitochondrial assays, we examined how clock disruption reprograms eCBome-associated lipid signaling and mitochondrial function. The objective of this study was to elucidate the molecular relationship between circadian rhythm disruption, eCBome remodeling, and mitochondrial dysfunction in hepatocytes, with particular focus on the DHEA-GPR110 signaling axis.

## Results

### Circadian Disruption Reduces Hepatic BMAL1 and GPR110 Expression and Alters Mitochondrial Dynamics

This study builds upon our previous observations demonstrating that circadian rhythm disruption alters eCBome profiles. In our earlier study, circadian rhythm was disrupted by keep the mice in constant light for 15 days and circadian disruption in mouse liver was associated with significantly reduced levels of eCBome NAEs DHEA and LEA, alongside increased eCBome MAG levels, including 2-OG and 2-SDG, as well as elevated plasma cytokines including IL-1β, IL-13, GM-CSF, and IFN-γ (43). To further investigate hepatic alterations associated with circadian disruption, we re-examined liver samples from the *in vivo* study. Hepatic BMAL1 protein levels were significantly reduced at CT11 in LL-exposed mice, suggesting the possible disruption of the liver circadian rhythm (Fig. 1A). This observation is consistent with previous studies reporting that constant light exposure disrupts molecular clock function and alters BMAL1 expression (4). While our single time-point measurement does not allow assessment of circadian phase, amplitude, or rhythmicity, it demonstrates a reduction in hepatic BMAL1 abundance at CT11 following LL exposure. We then went on to assess effects on the expression of eCBome genes at this time point using a custom qPCR array in male mice and preliminary results showed significant downregulation in GPR110 (supplementary figure1). Confirmation of the reduction in expression of GPR110 was then carried out by individual qPCRs in male as well as female mice. Given the emerging relationship between circadian regulation, eCBome signaling, and mitochondrial function, we next evaluated key genes involved in mitochondrial dynamics by assessing the expression of FIS1, OPAR, MFN1 and MFN2. Only fission-related gene FIS1 expression was found to change, being significantly increased under LL (Fig. 1B). We further assessed mitochondrial abundance by quantifying mitochondrial copy number and observed no significant differences in liver tissue (Fig. 1C).

**Figure 1.**
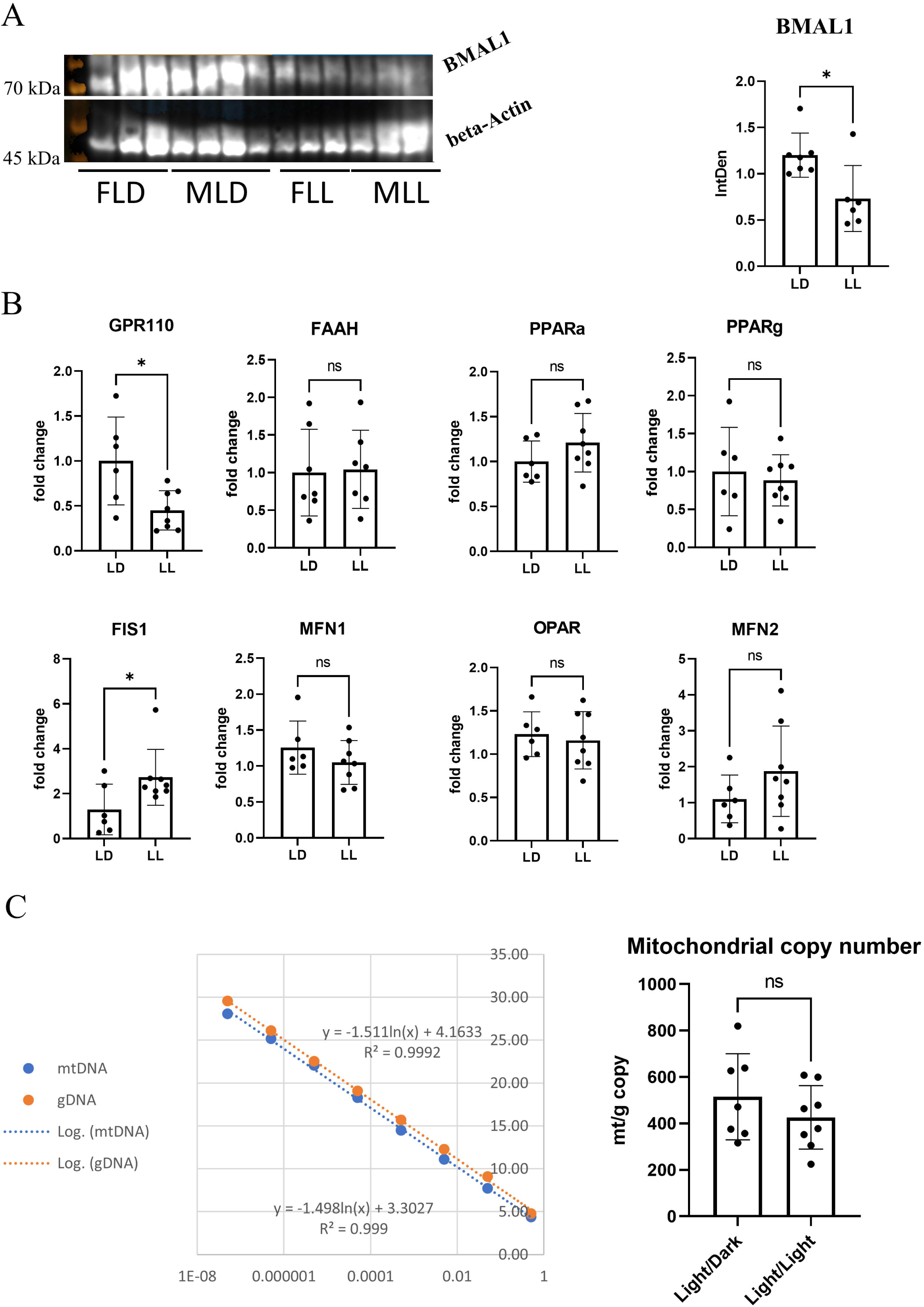
(A) Circadian disruption reduces hepatic BMAL1 protein expression. Representative western blot and densitometric quantification of BMAL1 protein expression in liver samples from mice maintained under a standard light/dark cycle (LD; 12 h light:12 h dark; n = 7) or exposed to constant light conditions (LL; 24 h light:0 h dark; n = 6). BMAL1 protein levels were normalized to β-actin and quantified by densitometric analysis. Data are presented as mean ± SD. p < 0.05 versus LD. **(B) Relative mRNA expression of genes involved in endocannabinoid metabolism and inflammatory signaling** (Daglab, Faah, GPR110, Pparg, Ppara, Mgll) and mitochondrial dynamics (FIS1, MFN2, OPA1, and MFN1) in liver samples from mice maintained under LD (n = 7) or LL (n = 8) conditions. Gene expression was quantified by qPCR and expressed as fold change relative to the LD group after normalization to housekeeping genes. Data are presented as mean ± SEM. p < 0.05 versus LD; ns, not significant. **(C) Mitochondrial DNA (mtDNA) copy number analysis in liver samples from mice maintained under LD (n = 7) or LL (n = 6) conditions.** Total DNA was extracted and quantitative PCR was performed using mitochondrial-specific primers and the genomic reference gene B2M. Standard curves generated from serially diluted PCR products were used to quantify mtDNA and genomic DNA (gDNA). Mitochondrial copy number was calculated as the mtDNA/gDNA ratio. Representative standard curves are shown on the left and quantified mtDNA copy number on the right. Data are presented as mean ± SEM. ns, not significant.

To investigate the role of BMAL1 in regulating hepatic lipid metabolism and inflammatory signaling, we generated BMAL1 knockout (KO) HepG2 cells using CRISPR/Cas9. This model allowed us to assess the cellular consequences of BMAL1 deficiency and its potential contribution to the alterations observed *in vivo* (43). Two independent clones (KO1 and KO2) were selected, and sequencing validated the successful editing of the target region, as did western blot analysis at the protein level (Fig. 1A). To compare the *in vitro* model with our previous *in vivo* findings, HepG2 control and BMAL1 KO cells were cultured in parallel and subjected to targeted lipidomics, TaqMan array gene expression analysis of eCBome associated and inflammatory genes, and cytokine profiling. Targeted lipidomics revealed that several fatty acids (e.g., docosahexaenoic acid (DHA), arachidonic acid (AA), and linoleic acid (LA)), together with downstream prostanoids (e.g., prostaglandin E2 (PGE2), prostaglandin F2α (PGF2α), and thromboxane B2 (TXB2)) and MAGs (e.g., 2-linoleoylglycerol (2-LG), 2-docosahexaenoylglycerol (2-DHG), and 2-AG), were significantly increased in BMAL1 knockout cells. In contrast, NAEs, including *N*-oleoyl-ethanolamine (OEA), and palmitoyl-ethanolamide (PEA), were significantly decreased, although *N*-eicosapentaenoyl-ethanolamide (EPEA) showed an opposite pattern. Overall, the *in vitro* model showed trends similar to those observed *in vivo*, with MAGs generally increased and NAEs decreased. Interestingly, within the MAG family, 2-OG deviated from this pattern and was significantly reduced in the knockout models, while this metabolite was significantly increased in the liver of mice on a disrupted cycle (43) (Fig. 2B). Because BMAL1 KO1 and BMAL1 KO2 displayed similar lipid profiles, one colony (BMAL1 KO1 cells) was selected for subsequent gene expression and cytokine analyses. Gene expression analysis revealed significant upregulation of GPR110 together with coordinated trend of activation of inflammatory pathways, including increased expression of CCL27 and GIPR in the KO model (Fig. 2C). Increased GPR110 expression was further validated by immunocytochemistry (ICC) at the protein level (Fig. 2D). Consistent with the elevated fatty acid profile, AdipoRed analysis demonstrated significantly increased total lipid accumulation in BMAL1 KO cells (Fig. 2E). Having elevated proinflammatory fatty acids and the MAGs encouraged us to profile cytokines. We found that pro-inflammatory cytokines including IL-6, IL-8, and TNF-α, were markedly increased (Fig. 2F), whereas IFN-γ and IL-12p70 were reduced in HepG2 BMAL1 KO cells.

**Figure 2.**
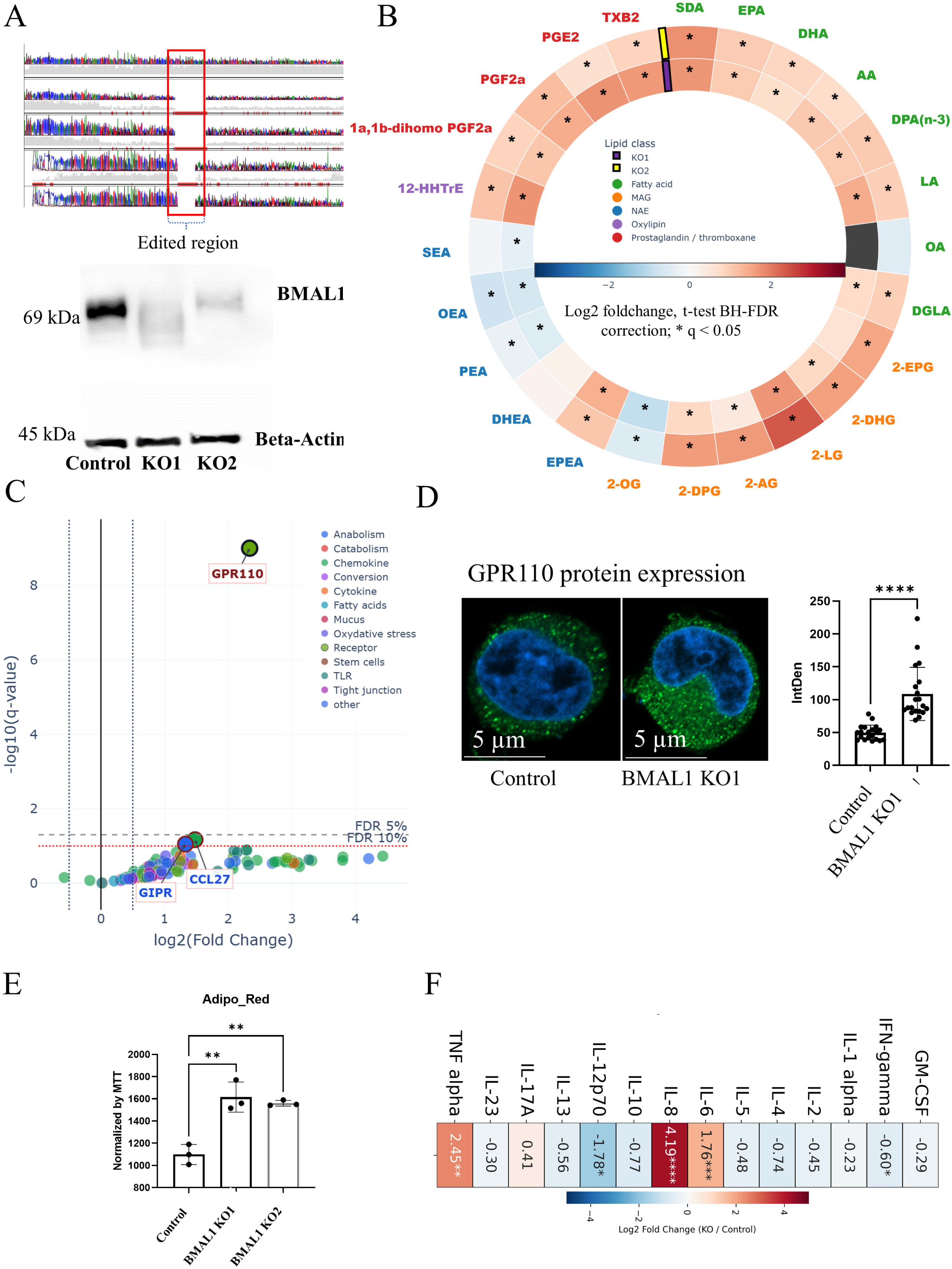
(A) Generation and validation of BMAL1 knockout (KO) HepG2 cells using the CRISPR/Cas9 system as an *in vitro* model of circadian disruption. Gene editing was validated by sequencing analysis, with the edited region highlighted (top). BMAL1 protein expression as assessed by western blot analysis in two independent BMAL1 KO clones and compared with control cells. β-actin was used as a loading control for normalization. **(B) Circular heatmap visualization of lipid mediator alterations across experimental groups.** Log2 fold changes relative to Control are shown for lipid mediators grouped into functional classes, including *N*-acylethanolamines (NAEs), monoacylglycerols (MAGs), fatty acids, prostaglandins/thromboxanes, and oxylipins. Red indicates increased abundance and blue indicates decreased abundance relative to Control. Comparisons included BMAL1 knockout clones (KO1 and KO2) versus Control. Statistical significance was assessed using Welch’s t-test followed by Benjamini–Hochberg false discovery rate (FDR) correction. Lipid mediators passing an FDR threshold of q < 0.1 were included in the heatmap, while those reaching q < 0.05 were considered statistically significant and are indicated by an asterisk (*). **(C) Volcano plot of differential qPCR gene expression profiles.** Differential expression was calculated using ΔCt/ΔΔCt-derived log2 fold changes of BMAL1KO1 cells relative to control samples. Statistical comparisons were performed using Welch’s t-test with n=3 per condition followed by Benjamini–Hochberg FDR correction. Significant genes (q ≤ 0.10) are labeled. Dashed vertical lines indicate log2FC thresholds (±0.5), while horizontal lines represent FDR thresholds (q = 0.10 and q = 0.05). The above gray line, considered as significant (q ≤ 0.05) and under gray line labeled genes consider trend. **(D) Representative confocal immunofluorescence images showing GPR110 expression in control and BMAL1 KO1 cell lines.** Nuclei were counterstained with Hoechst. Images were acquired using a confocal microscope under a 63× objective lens. Adjacent graph represents quantification of GPR110 fluorescence intensity measured as integrated density per cell using ImageJ software. Data are presented as mean ± SEM and analyzed using Student’s t-test. P < 0.05 was considered statistically significant. **(E) Lipid accumulation was assessed in control and BMAL1 knockout (KO) HepG2 cells using the AdipoRed assay**. Measurements were performed in control cells and two independent BMAL1 KO clones (KO1 and KO2) and normalized to MTT values to account for differences in cell number and viability. Data are presented as mean ± SEM. Statistical analysis was performed using one-way ANOVA followed by Tukey’s post hoc test. Individual data points are shown, and asterisks indicate statistically significant differences (*p < 0.05, ** p < 0.01, *** p < 0.001). **(F) Cytokine profiling in control and Bmal1 knockout (KO) HepG2 cells. Cytokine analysis was performed using one BMAL1 KO clone (KO1) and control cells (n = 3 per group).** Cytokine levels were quantified and expressed as log2 fold change relative to control cells. Statistical analysis was performed using one-way ANOVA followed by Tukey’s post hoc test on log2-transformed data. Red indicates increased cytokine levels and blue indicates decreased cytokine levels in BMAL1 KO cells relative to controls. Asterisks indicate statistically significant differences (p < 0.05).

### BMAL1 deletion disrupts circadian rhythmicity of DHEA and enhanced MAGs level

To induce circadian synchronization, cells were subjected to a 2 hours serum shock using 50% FBS followed by replacement with standard culture medium containing 10% FBS (44, 45). Circadian rhythmicity was then assessed using the pLV6-Bmal-luc luciferase reporter construct, which contains BMAL1 regulatory elements upstream of the luciferase gene. Because the BMAL1 promoter includes regions responsive to circadian transcription factors such as ROR and REV-ERB, synchronized cells were expected to exhibit oscillatory luciferase activity with progressively dampened amplitude over time (46, 47). Consistent with this expectation, following 50% FBS synchronization, control cells displayed rhythmic oscillations in promoter activity, indicating activation of the BMAL1 promoter and induced dynamic regulation of the circadian transcriptional network involving REV-ERB and ROR elements (Fig. 3A). In contrast, BMAL1 KO cells exhibited a lack of rhythmic oscillation and showed only baseline promoter activity following 50% FBS stimulation, suggesting impaired REV-ERB/ROR-mediated transcriptional feedback and disruption of the regulatory mechanisms required for normal circadian rhythmicity due to lack of BMAL1 protein. In parallel, lipid extracts were collected at multiple time points (including, 0 hour (unsynchronized), 2 hours (this samples were collected write after 2 hours of serum shock), 6 hours, 12 hours, and 24 hours) to determine whether circadian disruption altered temporal lipid profiles. Most NAEs (e.g., PEA, OEA, LEA, AEA), exhibited rhythmicity in both control and BMAL1 KO cells, although oscillation amplitudes were reduced in the knockout model. In contrast, DHEA completely lost rhythmicity in BMAL1 KO cells. Among MAGs, 2-LG, 2-AG, 2-EPG, 2-DPG, and 2-DHG maintained rhythmic oscillation in both control and knockout cells, whereas 2-OG lacked significant rhythmicity under either condition. Although 2-OG levels were lower at the unsynchronized baseline (0 hours), later it responded to serum shock similarly to other MAG molecules and showed increased levels in BMAL1 KO cells following synchronization. In addition, with the exception of OA, free fatty acid levels were generally elevated in BMAL1 KO cells (Fig. 3B). The persistence of rhythmicity in several lipid mediators despite BMAL1 deletion suggests that disruption of a core circadian component may not completely abolish oscillatory behavior. Residual rhythmic regulation may be maintained through other clock-associated factors, including REV-ERB and ROR pathways, which could partially compensate and preserve temporal patterns in downstream metabolic processes (48). Given the significant alterations in DHEA levels and its receptor GPR110 observed in both our *in vivo* and *in vitro* models, together with previous reports demonstrating that GPR110 is also implicated in fatty liver progression (49, 50). We hypothesized that DHEA-GPR110 signaling may provide a mechanistic link between circadian rhythm regulation and the endocannabinoidome.

**Figure 3.**
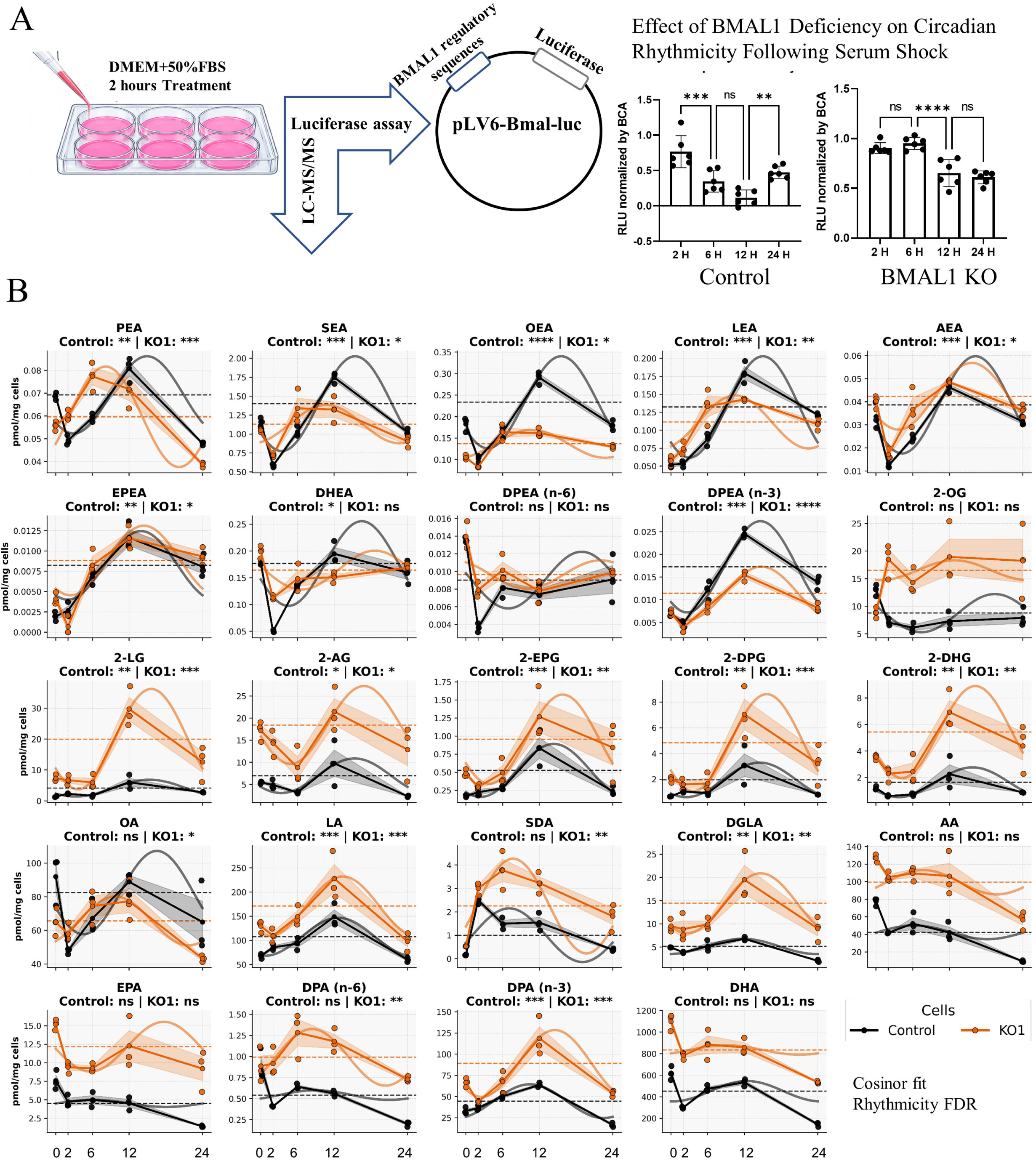
(A) Validation of serum shock-induced circadian rhythm synchronization in HepG2 control and BMAL1KO cells using a BMAL1 promoter luciferase reporter assay. HepG2 cells stably expressing the pLV6-Bmal-luc construct, containing BMAL1 regulatory sequences upstream of the luciferase reporter gene, were exposed to 50% fetal bovine serum (FBS) for 2 hours to induce circadian synchronization. Luciferase activity was measured at 2, 6, 12, and 24 h following serum shock to assess temporal changes in BMAL1 promoter activity. Protein extracts were collected at each time point and analyzed using a luciferase assay. Relative light units (RLU) were normalized to total protein content measured by the bicinchoninic acid (BCA) assay. Data are presented as mean ± SD. Statistical analysis was performed using one-way analysis of variance (ANOVA) followed by Tukey’s post hoc test. Individual data points are shown, and asterisks indicate statistically significant differences ((*p < 0.05, ** p < 0.01, *** p < 0.001). **(B) Time-dependent rhythmic profiling of lipid mediators in control and BMAL1 knockout (KO1) HepG2 cells.** Cells were synchronized by 50% FBS serum shock, and lipid levels were measured at multiple time points in control and BMAL1 KO1 cells (n = 3 per time point). Lipid concentrations were quantified by targeted liquid chromatography–tandem mass spectrometry (LC-MS/MS) and expressed as pmol/mg cells. Data are presented as mean ± SD with individual data points shown. Rhythmicity was evaluated using 24 hours cosinor analysis, and rhythmic significance was determined following false discovery rate (FDR) correction. Solid curves represent fitted cosinor models and dashed lines indicate the MESOR (midline estimating statistic of rhythm). Rhythmicity significance for each condition is indicated above each panel, where * indicates FDR-adjusted q < 0.05, ** indicates q < 0.01, *** indicates q < 0.001, and **** indicates q < 0.0001. Amplitude changes in BMAL1 KO1 cells relative to control are annotated within each plot.

### DHEA Alters Inflammatory, Lipid Metabolic, and eCBome Regulatory Networks

We next investigated the biological effects of DHEA on eCBome. HepG2 cells were treated with increasing concentrations of DHEA to identify an effective dose for subsequent experiments based on its previously reported effects on lipid metabolism in HepG2 cells (25). DHEA treatment significantly reduced total lipid levels in a dose-dependent manner (Fig. 4A). Both 10 and 100 nM DHEA elicited measurable biological responses and 100 nM was selected for subsequent experiments. To reproduce previously reported lipid-lowering effects of DHEA, cells in this initial experiment were co-treated with oleic acid (OA) to induce lipid accumulation.

**Figure 4.**
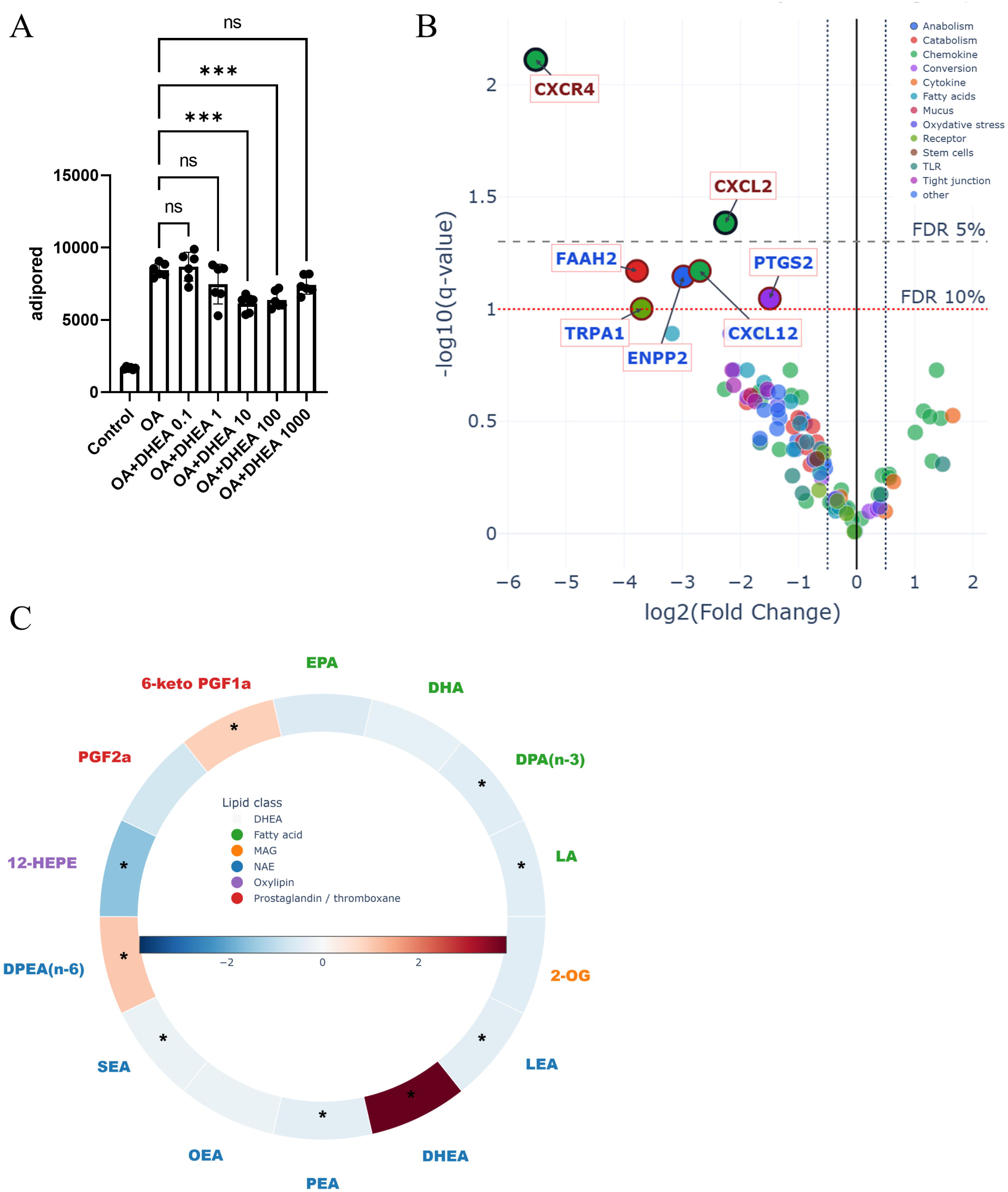
(A) Lipid accumulation in HepG2 cells following docosahexaenoyl ethanolamide (DHEA) treatment in the presence of oleic acid (OA) was assessed using the AdipoRed assay. Cells were treated with increasing concentrations of DHEA in the presence of OA, and total lipid accumulation was quantified (n = 5 per group). Data revealed a concentration-dependent, non-linear response to DHEA treatment. Data are presented as mean ± SD with individual data points shown. Statistical analysis was performed using one-way analysis of variance (ANOVA) followed by Tukey’s post hoc test. Asterisks indicate statistically significant differences (*p < 0.05, ** p < 0.01, *** p < 0.001); ns, not significant. **(B) Volcano plot of differential qPCR gene expression profiles.** Differential expression was calculated using ΔCt/ΔΔCt-derived log2 fold changes of DHEA treatment relative to control samples. Statistical comparisons were performed using Welch’s t-test with n=3 per condition followed by Benjamini–Hochberg FDR correction. Dashed vertical lines indicate log2FC thresholds (±0.5), while horizontal lines represent FDR thresholds (q = 0.10 and q = 0.05). The above gray line labeled genes, considered as significant (q ≤ 0.05) and under gray line labeled genes consider trend. **(C) Circular heatmap visualization of lipid mediator alterations in response to DHEA treatment for 24 hours.** Log2 fold changes relative to Control are shown for lipid mediators grouped into functional classes, including *N*-acylethanolamines (NAEs), monoacylglycerols (MAGs), fatty acids, prostaglandins/thromboxanes, and oxylipins. Red indicates increased abundance and blue indicates decreased abundance relative to Control. Statistical significance was assessed using Welch’s t-test followed by Benjamini–Hochberg false discovery rate (FDR) correction. Lipid mediators passing an FDR threshold of q < 0.1 were included in the heatmap, while those reaching q < 0.05 were considered statistically significant and are indicated by an asterisk (*).

However, OA was not used in downstream experiments because of its known effects on endocannabinoid-related pathways, including modulation of NAPE-PLD and FAAH activity (51). In addition, OA serves as a precursor for OEA production and can directly influence PPAR-α signaling (52), potentially confounding interpretation of DHEA-specific effects.

We next investigated whether DHEA directly modulates eCBome and inflammation pathways *in vitro*. HepG2 cells were treated with 100 nM DHEA and transcriptomic and lipidomic responses were assessed. DHEA treatment resulted in broad downregulation of genes involved in inflammatory signaling, including CXCL2, CXCR4 as well in a trend of decrease toward decrease in TRPA1, ENPP2, FAAH2, CXCL12, and PTGS2 (Fig. 4B). Lipidomic analysis revealed marked reductions in multiple eCBome-related lipids following DHEA treatment, including 12-HEPE, several NAEs (LEA, PEA, and SEA), as well as LA and DPA(n-3), while DPEA(n-6) and 6-keto-PGF2α were significantly increased (Fig. 4C).

### DHEA-GPR110 signaling regulates Ca²^+^ homeostasis and mitochondrial bioenergetics

BMAL1 deficiency has been associated with marked mitochondrial structural and functional alterations, including mitochondrial swelling and reduced ATP production (11). In addition, mitochondrial dysfunction is a common feature of fatty liver pathogenesis (15, 53). Given the central role of mitochondria in cellular energy production and lipid metabolism, we next investigated mitochondrial bioenergetics (54). Since mitochondrial function is tightly linked to Ca²^+^ signaling (55), mitochondrial Ca²^+^ levels were also assessed.

We first went on to assess the effects of DHEA on mitochondrial activity by Seahorse mito-stress test. DHEA treatment (100 nM for 2 hours) enhanced multiple metabolic parameters (supplementary Fig. 1B for OCR and ECAR curves), including glycolytic and mitochondrial ATP production, basal glycolysis, proton efflux rate (PER) (Fig. 5A). In contrast, BMAL1 KO1 and BMAL1 KO2 cells exhibited depressed OCR and ECAR curves (supplementary Fig 1C for OCR and ECAR curves). The similar behaviour of both knockout cell lines in the seahorse assays was expected given their similarity in the evaluations carried out above. Interestingly, despite increased GPR110 expression, DHEA treatment failed to restore or influence any of the metabolic parameters quantified in the seahorse assay for either of the BMAL1 knock out cell lines (Fig. 5A).

**Figure 5.**
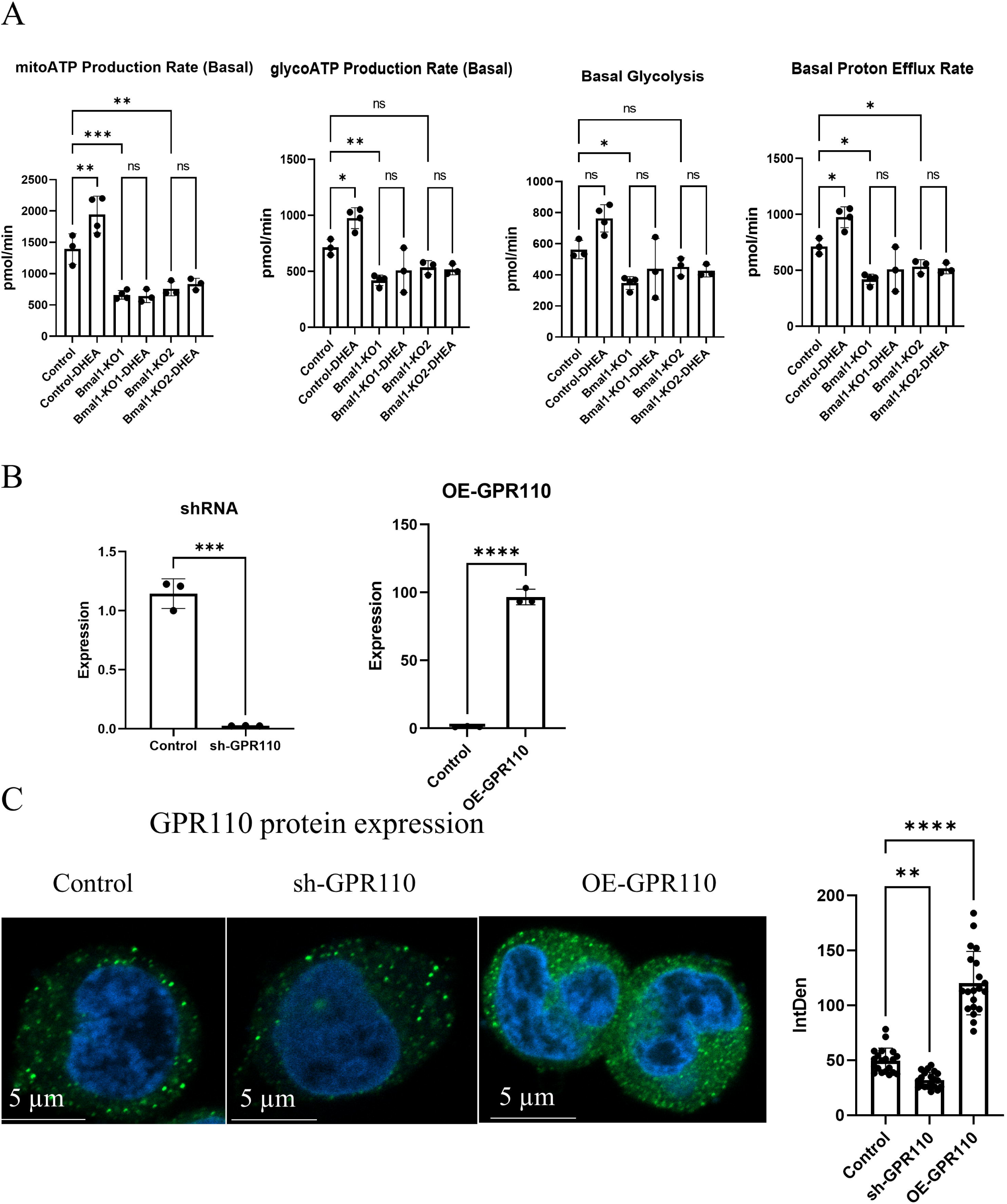
(A) Mitochondrial bioenergetic profiling of control and BMAL1 knockout (KO) HepG2 cells in response to docosahexaenoyl ethanolamide (DHEA) treatment using the Seahorse XF Mito Stress Test. Bioenergetic parameters including mitochondrial ATP (mitoATP) production rate, glycolytic ATP (glycoATP) production rate, basal proton efflux rate and basal glycolysis were assessed in control and BMAL1 KO clones (KO1 and KO2) with or without DHEA treatment. Data are presented as pmol/min with individual data points shown. Statistical analysis was performed using one-way analysis of variance (ANOVA) followed by Tukey’s post hoc test. Data are presented as mean ± SEM. Asterisks indicate statistically significant differences (*p < 0.05, ** p < 0.01, *** p < 0.001); ns, not significant**. (B) Validation of GPR110 knockdown (sh-GPR110) and overexpression (OE-GPR110) in HepG2 cells.** For gene silencing, cells were transduced with lentiviral vectors expressing short hairpin RNA (shRNA) targeting Gpr110 (shRNA pool contains 5 shRNA). For ectopic expression studies, cells were transduced with lentiviral vectors encoding the full-length GPR110 gene. The control sample is a mix of non-targeting scrambled control (shSCR) and pLX304 empty vector. Gene expression levels were quantified by quantitative PCR (qPCR) to validate knockdown and overexpression efficiency prior to functional experiments. Data are presented as mean ± SD with individual data points shown. Statistical analysis was performed using unpaired t-tests. Asterisks indicate statistically significant differences (p < 0.01). **(C) Representative confocal immunofluorescence images validating GPR110 expression in control and GPR110-manipulated HepG2 cells (Overexpression and knocked down).** Cells were stained with anti-GPR110 antibody and nuclei were counterstained with Hoechst. Images were acquired at 63× magnification using confocal microscopy. Quantification of GPR110 fluorescence intensity was performed using integrated density measurements per cell in ImageJ on 20 cells per group. Data are presented as mean ± SD. Statistical analysis was performed using Student’s t-test or one-way ANOVA as appropriate.

In order to determine whether the DHEA-induced metabolic effects observed above involved GPR110 signaling, we generated shRNA-mediated knockdown and ectopic overexpression of GPR110, respectively sh-GPR110, and OE-GPR110 HepG2 cells. Successful knockdown and overexpression of GPR110 was show at the RNA level by qPCR (Fig 5B) and protein level by Immunocytochemistry (ICC) (Fig. 5C). We then went on to examine the effects of DHEA on mitochondrial activity in manipulated GPR110 cell models.

sh-GPR110 (GPR110 knockdown) largely abolished the metabolic effects of DHEA. This suggests that DHEA effects on mitochondria are largely mediated by GPR110. Interestingly, OE-GPR110 (overexpression) significantly reduced glycolytic ATP production and proton efflux rate, whereas mitochondrial ATP production remained reactive to DHEA treatment and produced ATP (Fig. 6).

**Figure 6.**
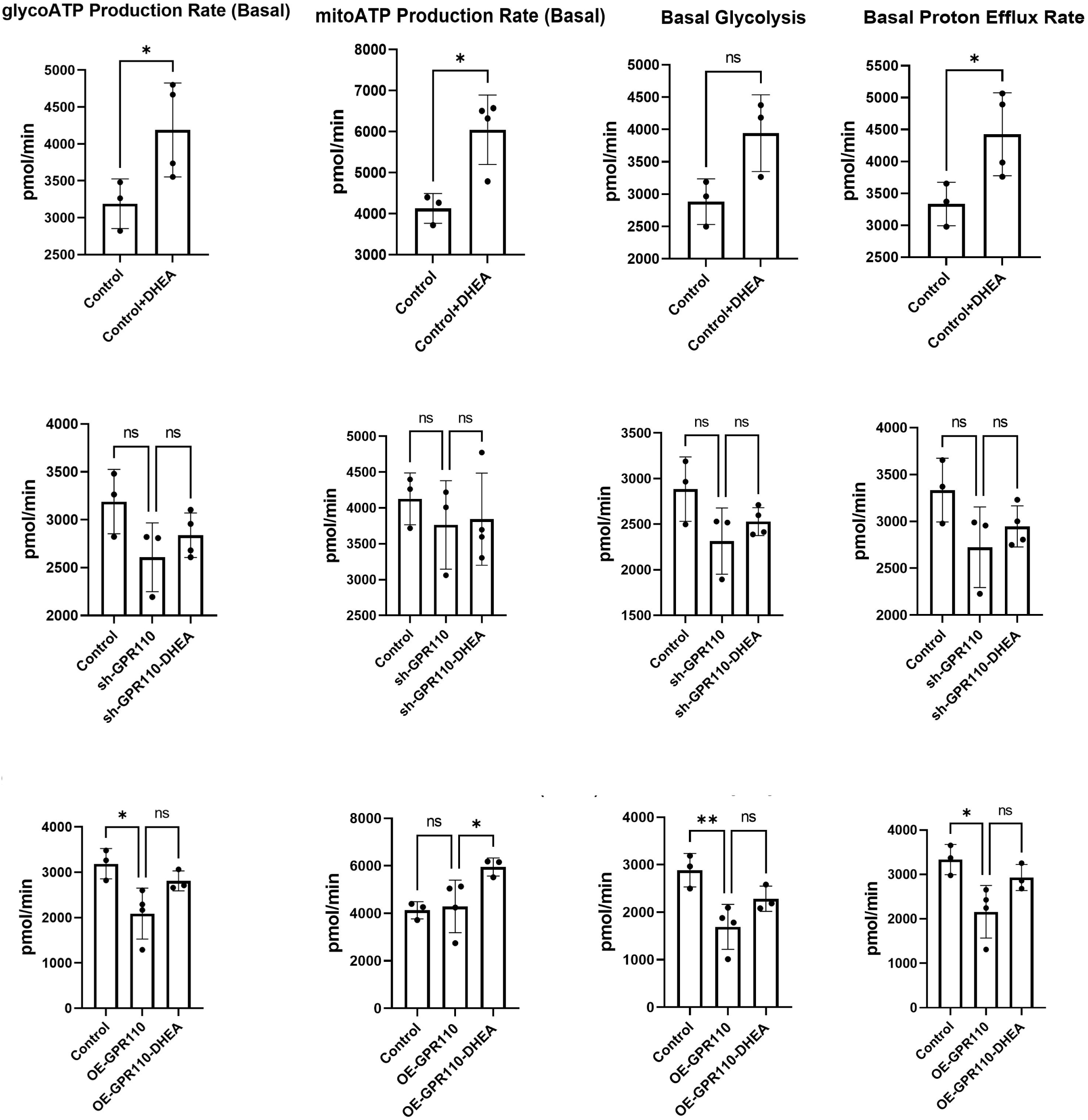
Mitochondrial bioenergetic profiling to investigate the role of GPR110 in mediating the effects of docosahexaenoyl ethanolamide (DHEA) in HepG2 cells using the Seahorse XF Mito Stress Test. Bioenergetic parameters including glycolytic ATP (glycoATP) production rate, mitochondrial ATP (mitoATP) production rate, basal glycolysis and basal proton efflux rate were assessed in control, GPR110 knockdown (sh-GPR110), and GPR110 overexpressing (OE-GPR110) HepG2 cells in the presence or absence of DHEA treatment. The control sample is a mix of non-targeting scrambled control (shSCR) and pLX304 (empty vector). Gene expression levels were quantified by quantitative PCR (qPCR) to validate knockdown and overexpression efficiency prior to functional experiments. Data are presented as pmol/min with individual data points shown. Statistical analysis was performed using one-way analysis of variance (ANOVA) followed by Tukey’s post hoc test. Data are presented as mean ± SD. Asterisks indicate statistically significant differences (*p < 0.05, ** p < 0.01, *** p < 0.001); ns, not significant.

To further investigate the effects of DHEA-GPR110 signalling on mitochondria, we employed mitochondrial- and endoplasmic reticulum (ER) -specific Ca²^+^ biosensors to assess mitochondrial and ER Ca²^+^ levels in real-time by flow cytometry (55).

The stable cell line containing the mitochondrial and ER biosensor was first validated with confocal microscope (Fig. 7A). Later, the signal intensity was measured in flowcytometry, we observed in control cell a spectrum of Ca²^+^ levels in mitochondria and ER, a feature that was a miss in the BMAL1 KO (Fig. 7B). Interestingly, both BMAL1KO1 and BMAL1KO2 clones displayed elevated mitochondrial and ER Ca²^+^ levels relative (mean AlexaFlour was used for stat) (Fig. 7C). The mitochondrial Ca²^+^ result was confirmed using a plate-reader using fluorescence area scan normalized to MTT (Fig. 7C). Additionally, fluorescence microscopy observation validated the already known swollen mitochondrial phenotype in the BMAL1KO cell model that might be due to overload of Ca²^+^ and upcoming osmotic water into mitochondria (Fig. 7D) (11).

**Figure 7.**
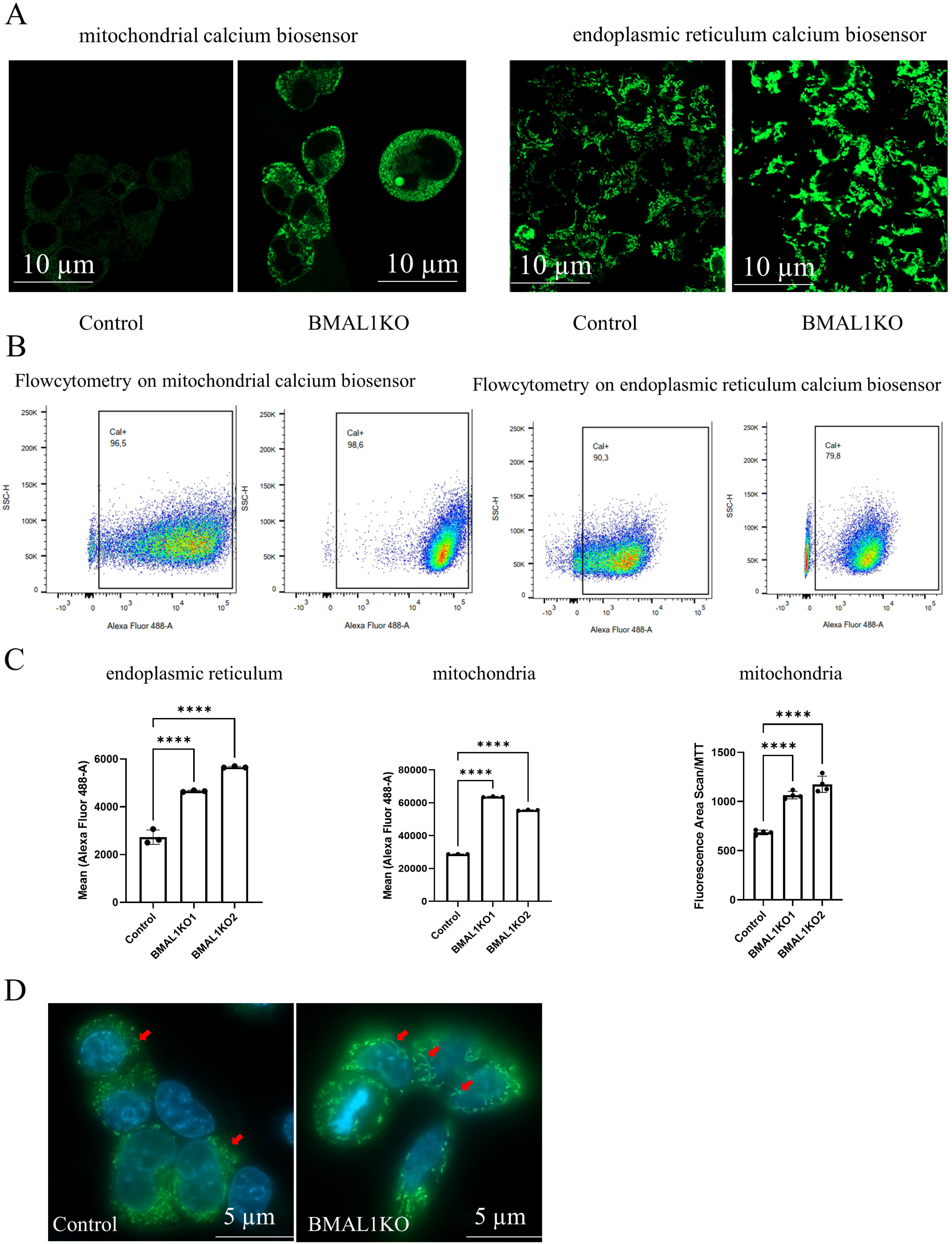
(A) Confocal validation of mitochondrial and ER Ca²^+^ biosensors in Control and BMAL1KO cells. Representative confocal microscopy images of stable cell lines expressing mitochondria- and ER-targeted Ca²^+^ biosensors. Fluorescence localization confirmed successful biosensor expression in both organelles and revealed increased signal intensity in BMAL1KO cells compared with controls, supporting subsequent Ca²^+^ analyses. **(B) Flow cytometric validation and analysis of mitochondrial and ER Ca²**^+^ **biosensor activity in Control and BMAL1KO cells.** Representative flow cytometry plots of Control and BMAL1KO stable cell lines expressing mitochondria- and ER-targeted Ca²^+^ biosensors. Ca²^+^-associated fluorescence was measured using Alexa Fluor 488-A, and Ca+ populations were identified through gating analysis. These data were used for quantification of organelle-specific Ca²^+^ alterations in BMAL1KO cells. **(C) BMAL1 deficiency increases ER and mitochondrial Ca²**^+^ **levels in HepG2 cells.** Ca²^+^levels in the endoplasmic reticulum (ER) and mitochondria were quantified in Control and BMAL1 knockout (KO1 and KO2) HepG2 cells expressing organelle-specific Ca²^+^ biosensors. ER and mitochondrial biosensor fluorescence was measured by flow cytometry (Alexa Fluor 488-A), and mitochondrial fluorescence area was additionally quantified and normalized to MTT. Data are presented as mean ± SD with individual data points shown. Statistical analysis was performed using one-way ANOVA followed by Tukey’s multiple comparison test. **** p < 0.05. **(D) BMAL1 deficiency is associated with mitochondrial swelling.** Representative fluorescence microscopy images demonstrate mitochondrial Ca²^+^(Green) and stained nuclei (Blue), enlarged and swollen mitochondrial structures in BMAL1KO cells (red arrows), consistent with previously reported mitochondrial abnormalities during circadian disruption.

Further experiments on DHEA-GPR110 models showed that DHEA treatment increased mitochondrial and ER Ca²□ levels within 2 hours of exposure (Fig. 8A). Accordingly, sh-GPR110 reduced mitochondrial and ER Ca²^+^ levels, whereas OE-GPR110 produced the opposite effect and significantly increased mitochondrial and ER Ca²^+^ level (Fig. 8A). Mitochondrial Ca²^+^ fluctuation was independently validated using a plate-reader fluorescence area scan normalized to MTT (Fig 8A). When the ability of DHEA to modify mitochondrial Ca²^+^ levels was tested in sh-GPR110 and OE-GPR110 cells its effects were lost in the former, but maintained in the latter. Together these data are inline with previous report on DHEA regulation of intracellular Ca²^+^ dynamics (56) and suggest that DHEA signals through GPR110 to increase mitochondrial Ca²^+^ levels which may account for increased in mitochondrial ATP production, and that constitutive over expression of GPR110 may lead to mitochondrial Ca²^+^ overload.

**Figure 8.**
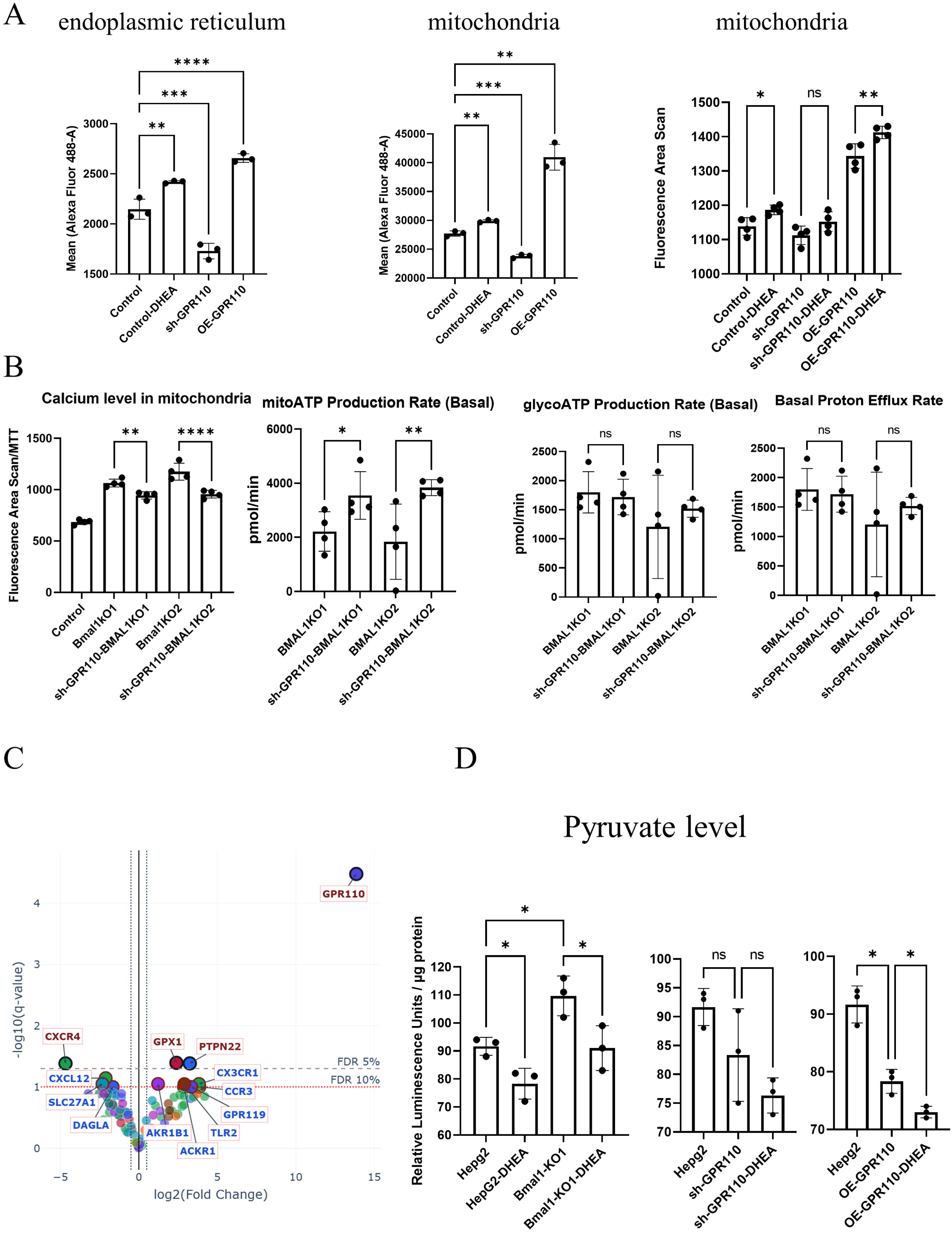
(A) GPR110 regulates mitochondrial Ca²^+^ homeostasis, bioenergetic function, inflammatory gene expression, and pyruvate metabolism in HepG2 cells. (A) Mitochondrial and endoplasmic reticulum (ER) Ca²^+^measurements were performed to investigate the role of GPR110 in mediating docosahexaenoyl ethanolamide (DHEA) responses in HepG2 cells. Mitochondrial Ca²^+^ measurements to investigate the role of GPR110 in mediating docosahexaenoyl ethanolamide (DHEA) responses in HepG2 cells. Mitochondrial Ca²+ levels were assessed by flow cytometry using Alexa Fluor 488 fluorescence in control, sh-GPR110, and OE-GPR110- HepG2 cells following DHEA treatment. The control sample is a mix of non-targeting scrambled control (shSCR) and pLX304 empty vector. ER (left panel) and mitochondrial (middle panel) Ca²^+^ levels were measured by flow cytometry and quantified using mean Alexa Fluor 488-A fluorescence intensity. Mitochondrial Ca²+ levels were additionally assessed independently in sh-GPR110 and OE-GPR110 cells using a plate-reader fluorescence area scan assay and quantified as fluorescence area values (right panel). Data are presented as mean ± SD with individual data points shown. Statistical analysis was performed using one-way ANOVA followed by Tukey’s multiple comparison test. *p < 0.05, **p < 0.01, ***p < 0.001; ns, not significant. (B) **Effects of GPR110 knockdown on mitochondrial Ca²**^+^ **levels and mitochondrial bioenergetics in BMAL1 knockout (KO) HepG2 cells.** BMAL1 KO1 and BMAL1 KO2 cells were transduced with a GPR110 short hairpin RNA (shRNA) pool to reduce GPR110 expression. Mitochondrial Ca²^+^ levels were quantified by fluorescence area scan normalized to MTT (left panel). Mitochondrial bioenergetic function was subsequently evaluated using the Seahorse XF Mito Stress Test, and mitochondrial ATP (mitoATP) production rate was quantified (middle and right panels). The middle panel compares control, BMAL1 KO, and Gpr110 knockdown conditions and was analyzed using one-way analysis of variance (ANOVA) followed by Tukey’s post hoc test, whereas the right panel was analyzed using unpaired t-tests. Data are presented as mean ± SD with individual data points shown. Asterisks indicate statistically significant differences (*p < 0.05, ** p < 0.01, *** p < 0.001); ns, not significant. **(C) Gene expression profiling of endocannabinoidome- and inflammation-related pathways in GPR110 overexpressing HepG2 cell model using a TaqMan gene expression array.** Volcano plot of differential qPCR gene expression profiles. Differential expression was calculated using ΔCt/ΔΔCt-derived log2 fold changes relative to control samples. Statistical comparisons were performed using Welch’s t-test with n=3 per condition followed by Benjamini–Hochberg FDR correction. Dashed vertical lines indicate log2FC thresholds (±0.5), while horizontal lines represent FDR thresholds (q = 0.10 and q = 0.05). The above gray line, considered as significant (q ≤ 0.05) and under gray line labeled genes consider trend. **(D) Pyruvate levels were quantified in control, GPR110-overexpressing, GPR110 knockdown, and BMAL1 knockout HepG2 cells with or without DHEA treatment using a luciferase-based pyruvate assay.** Luminescence values were normalized to total protein content measured by bicinchoninic acid (BCA) assay and expressed as relative luminescence units (RLU) per μg protein. Data are presented as mean ± SD with individual data points shown. Statistical analysis was performed using one-way ANOVA followed by Tukey’s multiple comparison test. *p < 0.05, **p < 0.01, ***p < 0.001; ns, not significant.

Later to further examine the relationship between GPR110 and mitochondrial Ca²^+^ regulation under circadian disruption, we used GPR110 shRNA to knock down GPR110 in BMAL1 KO cells. This manipulation reduced mitochondrial Ca²^+^ levels and was accompanied by increased mitochondrial ATP production in BMAL1 KO cells (normalized to MTT) (Fig. 8B).

The similarities between control cells with OE-GPR110 and BMAL1 KO cells in displaying elevated mitochondrial Ca²□ levels prompted further investigation into whether OE-GPR110 induces broader transcriptional adaptations. To address this, we employed the same TaqMan array used above to evaluate genes involved in eCBome signaling, inflammatory pathways, and metabolic regulation (Fig. 8C). OE-GPR110 induced significant increases in GPX1 and PTPN22 expression. Increased GPX1, an antioxidant enzyme involved in protection against oxidative stress, may reflect a compensatory response to altered mitochondrial activity and Ca²□ handling. Similarly, PTPN22, a regulator of immune and inflammatory signaling pathways, suggests modulation of cellular stress or inflammatory responses following GPR110 activation. In addition, several genes exhibited trends toward increased expression, including CX3CR1, CCR3, TLR2, and ACKR1, indicating potential remodeling of chemokine and immune-associated signaling pathways. Conversely, CXCR4 expression was reduced, accompanied by trends toward lower CXCL12, suggesting suppression of the CXCL12-CXCR4 signaling axis. Because this pathway contributes to inflammatory signaling, cellular migration, and metabolic regulation, the observed trends toward reduced SLC27A1 and DAGLA expression may further point toward altered fatty acid transport and endocannabinoid-related lipid metabolism. Collectively, these findings suggest that OE-GPR110 induces coordinated transcriptional changes involving oxidative stress responses, inflammatory signaling, and eCBome-associated metabolic pathways, potentially linking increased mitochondrial Ca²□ signaling with broader cellular adaptation mechanisms.

Given the observed effects of DHEA and GPR110 on mitochondrial bioenergetics and mitochondrial Ca²□ dynamics, we next investigated pyruvate metabolism. Pyruvate serves as a key metabolic intermediate linking glycolysis with mitochondrial oxidative metabolism through pyruvate dehydrogenase (PDH)-mediated entry into the tricarboxylic acid (TCA) cycle. Because mitochondrial Ca²□ can activate PDH and thereby facilitate pyruvate utilization and mitochondrial ATP production, we hypothesized that alterations in DHEA-GPR110 signaling may influence pyruvate handling.

Pyruvate measurements further supported the metabolic effects of DHEA (Fig. 8D). DHEA treatment significantly reduced pyruvate abundance in HepG2 cells, potentially reflecting enhanced mitochondrial utilization of pyruvate through increased oxidative metabolism. Consistent with this interpretation, DHEA also increased mitochondrial activity and mitochondrial Ca²□ levels in earlier experiments. OE-GPR110 overexpression similarly reduced pyruvate levels, suggesting that GPR110 signaling itself influences pyruvate metabolism. Additionally, DHEA treatment further decreased pyruvate levels in OE-GPR110 cells. In contrast, sh-GPR110 alone had little effect on basal pyruvate abundance but largely abolished the ability of DHEA to reduce pyruvate levels, further supporting a role for GPR110 in mediating DHEA responses.

Interestingly, BMAL1 KO cells exhibited elevated basal pyruvate levels relative to control cells. Increased pyruvate accumulation may indicate that glycolytic flux exceeds the capacity of mitochondria to further process pyruvate through oxidative metabolism, consistent with the impaired mitochondrial function observed in the knockout cells. This observation aligns with the reduced ATP production and elevated mitochondrial Ca²□ levels detected previously. Despite this, DHEA treatment significantly reduced pyruvate abundance in BMAL1 KO cells, suggesting that DHEA may partially restore pyruvate utilization despite circadian disruption. Together, these findings support a model in which DHEA-GPR110 signaling modulates mitochondrial Ca²□ dynamics and pyruvate utilization to regulate mitochondrial bioenergetics.

## Discussion

The present study suggests a mechanistic link between circadian disruption, eCBome remodeling, and Ca²□ homeostasis, mediated in part through DHEA-GPR110 signaling. By generating a BMAL1 KO HepG2 cell line, we recapitulated key molecular alterations previously observed *in vivo* under constant light exposure. BMAL1 deficiency induced substantial changes in bioactive lipid mediator profiles, intracellular Ca²□ dynamics, mitochondrial bioenergetics, and inflammatory signaling, collectively supporting that circadian transcriptional control may coordinate lipid signaling pathways with cellular energy metabolism.

The BMAL1 KO model demonstrated broad remodeling of fatty acids, prostaglandins, MAGs, and NAEs in unsynchronized cells, consistent with our earlier observations in liver tissue from mice exposed to constant light. The increase in MAG species, particularly 2-AG, 2-DHG, and 2-LG, may indicate enhanced endocannabinoidome signaling activity. Notably, MAGs, especially 2-AG, have been implicated in hepatic lipid accumulation and progression of NAFLD and alcoholic fatty liver through the activation of the cannabinoid receptor CB1 (57, 58). In contrast, reduced levels of saturated and monounsaturated NAEs, including PEA and OEA, which activate PPARA a key lipid metabolism-regulating transcription factor that is protective against NAFLD (35, 52) are frequently observed in liver disorders where their protective signaling functions are diminished (59, 60). BMAL1 KO hepatocytes also had increased levels of several free fatty acids (e.g., SDA, EPA, DHA and AA), together with elevations in prostaglandins, indicating extensive remodeling of hepatic lipid metabolism. Increased AA availability and prostaglandin production may reflect enhanced inflammatory signaling, whereas changes in ω-3 fatty acids such as SDA, EPA, and DHA may represent compensatory or adaptive responses given their ability to alleviate hepatosteatosis and inflammation (61). This interpretation is supported by the concomitant increase in multiple free fatty acids, prostaglandins, and MAG species observed in our study (62). Consistent with a pro-inflammatory environment, IL-8, IL-6 and TNF-alpha were significantly elevated in the circadian disruption model. Together, these findings suggest that eCBome alteration in BMAL1 is part of an adaptive mechanism or the consequence of proinflammatory pathways.

Among the genes altered by circadian disruption, GPR110 emerged as a potentially important mediator linking lipid signaling with intracellular Ca²□ regulation. Interestingly, unlike our *in vivo* mouse model in which hepatic GPR110 expression is reduced, GPR110 was significantly increased in the HepG2 BMAL1 KO model. Species-specific gene expression differences between mouse and human liver transcriptional programs have been extensively reported, with more than 50% of liver genes exhibiting opposite regulation patterns (63), including limited overlap in fatty liver disease signatures (64). Such differences may partly explain the distinct GPR110 responses observed between our *in vivo* and *in vitro* systems.

Interestingly, DHEA was the only NAE that lost rhythmicity in BMAL1 KO hepatocytes and it was already reported to have hepatic increased levels in the mouse model exposed to constant light (43). In contrast, although 2-OG levels were initially reduced under unsynchronized conditions, serum shock (50% FBS) induced an increase similar to that observed for other MAG species, suggesting that MAG regulation may depend more strongly on precursor availability. It is already shown that 2-OG significantly increases in response to constant light and circadian disruption (43) and in contrast to DHEA, which lowers the hepatocyte lipid accumulation (25), 2-OG enhances lipid accumulations (25, 65) and causes mitochondrial dysfunction (66). Previous studies have shown that in BMAL1 KO models, CLOCK, ROR, and Cry1 expression can remain rhythmic upon serum shock and, in some cases, become elevated (67). Therefore, one possible explanation for observing rhythmicity in several lipids in the BMAL1KO model is that serum shock induced rhythmic activity in these remaining clock components, which subsequently regulated downstream genes involved in lipid synthesis and metabolism. Interestingly, CRY1 (68), CLOCK and ROR directly regulate lipid metabolism (69, 70). This interpretation raises the possibility that genes responsible for MAG production (e.g. DAGLA/B) may be regulated primarily through CLOCK, ROR, or Cry1-dependent mechanisms rather than requiring direct BMAL1 control. Furthermore, DHEA and 2-OG displayed profiles distinct from other members of the NAE and MAG families, suggesting greater dependence on BMAL1 signaling. However, because the enzymes involved in NAE and MAG synthesis are not strictly substrate-specific (71, 72), the regulatory mechanisms underlying these selective effects likely remain to be defined.

The DHEA-GPR110 signaling pathway was conspicuously modified in both circadian rhythm’s models (mice and BMAL1KO cells). The consistent alteration of this pathway across both *in vivo* and *in vitro* models suggests that DHEA-GPR110 signaling may represent an important mechanism linking circadian disruption to metabolic remodeling. Consistent with this possibility, DHEA treatment broadly remodeled lipid mediator profiles, reducing 12-HEPE and several NAE species, including SEA, PEA and LEA, together with LA and DPA(n-3). Given that LA serves as a major fatty acid incorporated into cellular lipid pools and is the most prevalent omega-6 polyunsaturated fatty acid (PUFA) in the western diet (73), its reduction may partially be explained by the decrease in total lipid accumulation observed following DHEA treatment. In contrast, DHEA selectively increased levels of 6-keto-PGF1α, a stable metabolite of prostacyclin (PGI2) (74). 6-KETO PGF1 α is a marker for PGI2 production that negatively correlates with fatty liver disease progression severity of liver injury (75). Liver local production of PGI2 causes a vasodilatory and anti-thrombotic molecule which assists the flow of blood in the area and protects liver cells (76). This is particularly relevant in diabetes, where patient serum has been shown to impair PGI production in epithelial cells *in vitro* (77). These findings suggest that DHEA does not simply alter individual lipid mediators but induces broader remodeling of lipid metabolic networks. DHEA also altered the expression of genes associated with inflammatory signaling pathways, including CXCL2 and CXCR4. Because DHEA has been previously reported to exert anti-inflammatory actions (36), suppression of inflammatory mediators and chemokine-related signaling pathways may reflect a coordinated anti-inflammatory transcriptional response induced by DHEA in hepatocytes. Collectively, these findings suggest that exogenous DHEA induces adaptive remodeling of metabolic and inflammatory pathways.

Mitochondria function as central regulators of cellular energy production and are tightly controlled by circadian mechanisms that influence both their structure and function (78, 79). Mitochondrial dysfunction has been strongly associated with obesity and metabolic syndrome (80). Mitochondrial Ca²□ plays an essential role in bioenergetic regulation and its Ca²□ charge/discharge is necessary for mitochondrial ATP production (81). Under physiological conditions, increases in intracellular Ca²□ stimulate mitochondrial uptake, which enhances ATP production through activation of pyruvate dehydrogenase and key tricarboxylic acid cycle enzymes including IDH and α-KGDH (81–83). Following DHEA treatment, we observed increased mitochondrial and ER Ca²□ levels, which is in line with the previously reported DHEA regulation of Ca²□ entry into the cells (56). This increase in mitochondrial Ca²□ was accompanied by reduced pyruvate abundance and enhanced proton efflux rate, glycolytic and mitochondrial ATP production. Interestingly, growing evidence indicates that several glycolytic enzymes can localize to the outer mitochondrial membrane and form functional glycolytic complexes that facilitate direct coupling between glycolysis and oxidative phosphorylation (84). These observations suggest that DHEA may improve shuttling of pyruvate into the mitochondrial matrix to fuel ATP production under conditions in which glycolysis and proton efflux are increased.

Consistent with our earlier observations that the DHEA-GPR110 signaling pathway was conspicuously altered in *in vivo* and *in vitro* models, functional manipulation of GPR110 further supported its role in regulating organelle Ca²□ homeostasis and metabolic responses. Sh-GPR110 reduced mitochondrial and ER Ca²□ levels, whereas GPR110 overexpression (GPR110-OE) increased Ca²□ accumulation in both organelles. As expected, sh-GPR110 abolished responsiveness to DHEA, supporting the requirement of GPR110 for DHEA-associated effects. In contrast, OE-GPR110 reduced basal proton efflux rate, glycolysis, and glycolytic ATP production, showing a bioenergetic profile similar to BMAL1KO cells (which already exhibit elevated GPR110 expression), while preserving mitochondrial ATP responsiveness to DHEA. These findings suggest that DHEA may exert dual metabolic effects by enhancing both glycolytic and mitochondrial ATP production, whereas excessive Ca²□ accumulation induced by GPR110 overexpression selectively disrupts glycolysis-associated ATP generation while leaving mitochondrial ATP production relatively intact. This metabolic phenotype may resemble features reported in specific tumor types that do not fully conform to the classical Warburg effect. GPR110 has been reported to be overexpressed in several cancers including lung and prostate cancer (85), glioma (86), HER2-positive breast cancer (85), and gastric cancer (87). GPR110 has also been implicated in NAFLD and MASH progression (49, 50), both recognized risk factors for hepatocellular carcinoma (HCC)(87). Interestingly, several tumors exhibit greater dependence on oxidative phosphorylation (OXPHOS) rather than glycolysis. Early prostate adenocarcinoma, non-proliferative hepatocellular carcinoma, and liver cancer stem cells have all demonstrated substantial reliance on mitochondrial ATP production (88–92). Therefore, the preservation of mitochondrial ATP production despite impaired glycolytic ATP generation observed in the OE-GPR110 model may reflect a metabolic shift toward mitochondrial energy dependence similar to that reported in these tumor settings.

Considering that mitochondrial ATP production is tightly linked to Ca²□ homeostasis (charge/discharge) (83), physiological Ca²□ uptake stimulates mitochondrial metabolism, whereas sustained Ca²□ overload can induce mitochondrial permeability transition, resulting in mitochondrial swelling and impaired function (93). Elevated mitochondrial Ca²□ observed in BMAL1 KO cells may therefore contribute to the reduction in both glycolytic and mitochondrial ATP production. Importantly, the metabolic phenotype differed between BMAL1 KO and OE-GPR110 models. In OE-GPR110 cells, mitochondrial ATP production remained functional despite elevated Ca²□ levels, whereas in BMAL1 KO cells both mitochondrial and glycolytic ATP production were impaired. Supporting this interpretation, GPR110 knockdown in BMAL1 KO cells reduced mitochondrial Ca²□ accumulation while restoring mitochondrial ATP production, suggesting that excessive GPR110-associated Ca²□ loading contributes to mitochondrial dysfunction under circadian disruption. This observation is also consistent with the swollen mitochondrial morphology observed in BMAL1 KO cells. These findings suggest that DHEA influences cellular metabolism across multiple interconnected steps, extending from pyruvate utilization and glycolysis to mitochondrial bioenergetics. The reduction in pyruvate abundance together with increased glycolytic and mitochondrial ATP production following DHEA treatment may indicate enhanced pyruvate utilization within mitochondria to support oxidative metabolism. In contrast, although GPR110 overexpression preserved mitochondrial ATP production despite elevated mitochondrial Ca²□ levels, this effect may reflect maintenance of mitochondrial oxidative pathways, potentially through sustained activation of Ca²□ -sensitive enzymes involved in the tricarboxylic acid (TCA) cycle. In BMAL1 KO cells, however, excessive mitochondrial Ca²□ accumulation was associated with impaired mitochondrial and glycolytic ATP production, suggesting a distinct metabolic state. Under physiological conditions, mitochondrial Ca²□ enhances ATP generation through activation of pyruvate dehydrogenase and TCA cycle enzymes; however, sustained Ca²□ overload can become detrimental. One possible explanation is that prolonged mitochondrial Ca²□ accumulation under circadian disruption alters metabolic flux and compromises efficient substrate utilization, shifting mitochondria from a bioenergetic-supportive state toward dysfunction. This interpretation is consistent with the impaired ATP production and swollen mitochondrial morphology observed in BMAL1 KO cells.

## Conclusions

The present study identifies a potential mechanistic link between circadian disruption, endocannabinoidome remodeling, mitochondrial Ca²□ homeostasis, and inflammatory signaling in hepatocytes, with DHEA-GPR110 signaling emerging as an important mediator within this network. Loss of BMAL1 induced broad alterations in lipid metabolism characterized by increased MAGs, reduced levels of protective NAEs, enhanced inflammatory responses, and impaired mitochondrial bioenergetics. Our findings suggest that disruption of circadian regulation promotes a shift toward a pro-inflammatory and metabolically dysregulated state that includes alterations in bioactive eCBome lipids levels, associated with altered Ca²□ handling and reduced mitochondrial efficiency. Functional manipulation of the DHEA-GPR110 signalling pathway further indicated its involvement in regulating mitochondrial Ca²□ dynamics and ATP production, with DHEA exhibiting protective effects by reducing lipid accumulation and modulating inflammatory and metabolic pathways. Collectively, these findings support a model in which circadian regulation contributes to hepatic metabolic homeostasis through coordinated interactions between bioactive lipid signaling and mitochondrial function.

## Material and methods

### Plasmids and Antibodies

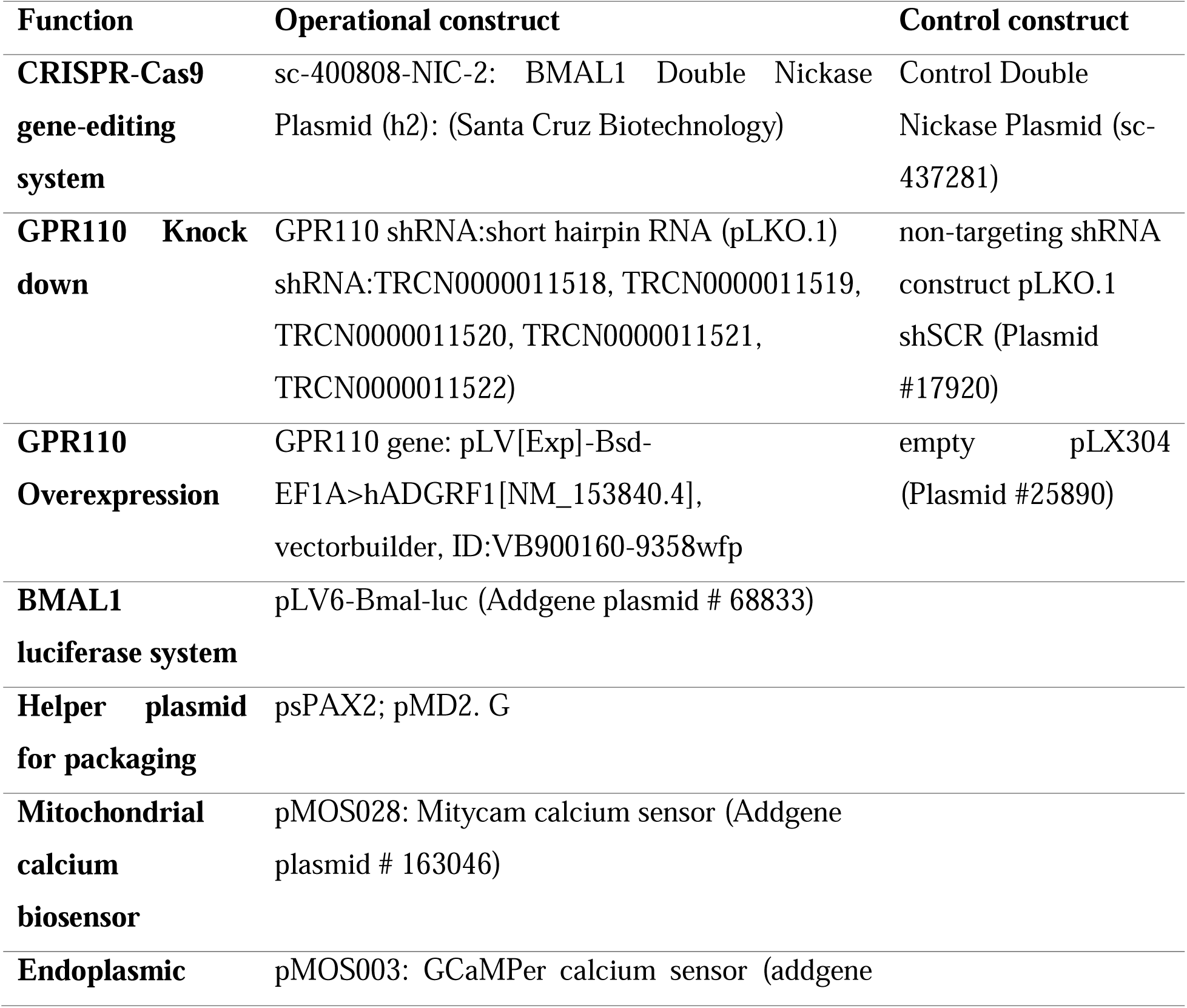
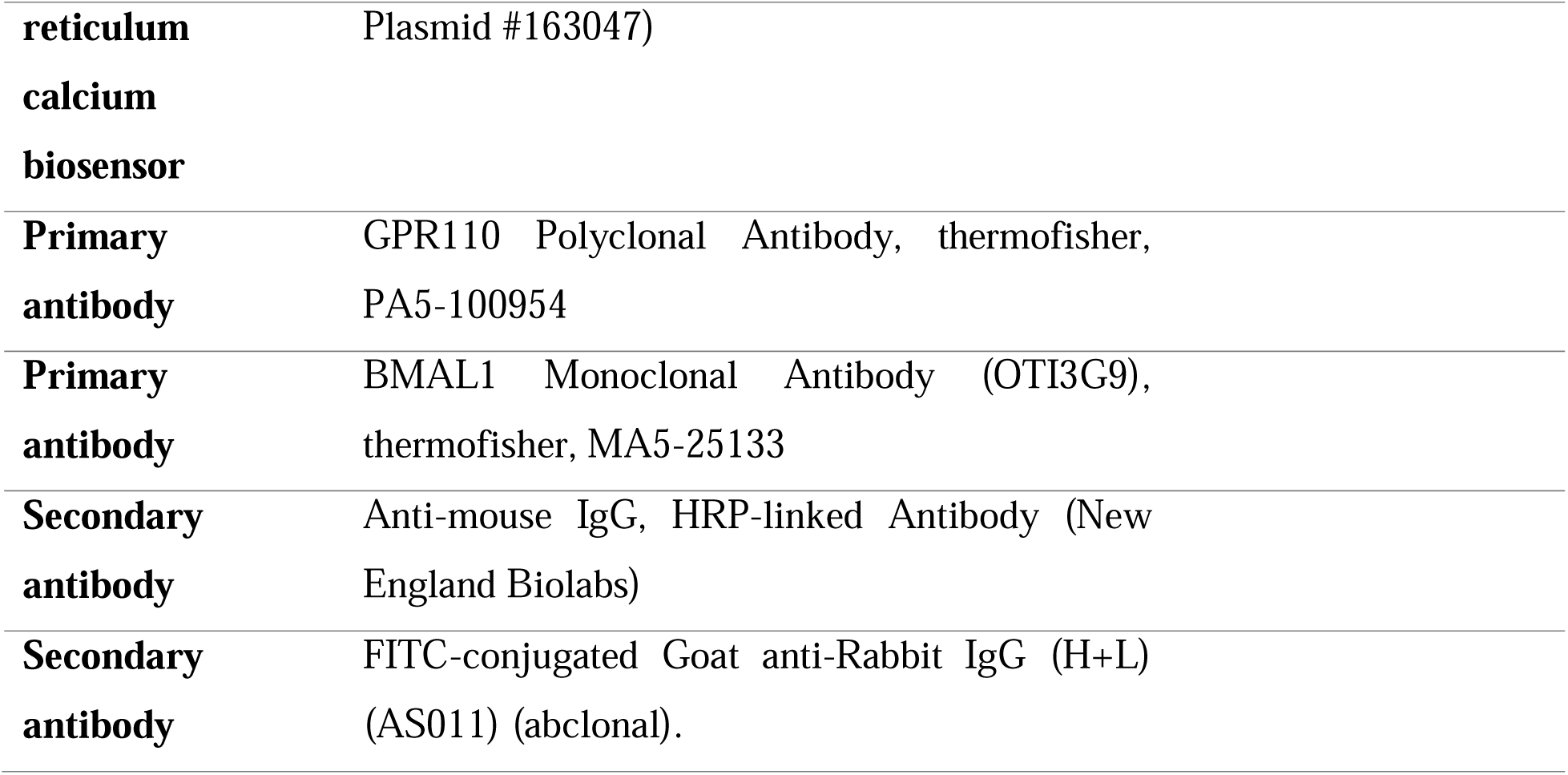

| Function | Operational construct | Control construct |
| --- | --- | --- |
| **CRISPR-Cas9 gene-editing system** | sc-400808-NIC-2: BMAL1 Double Nickase Plasmid (h2): (Santa Cruz Biotechnology) | Control Double Nickase Plasmid (sc-437281) |
| **GPR110 Knock down** | GPR110 shRNA:short hairpin RNA (pLKO.1) shRNA:TRCN0000011518, TRCN0000011519, TRCN0000011520, TRCN0000011521, TRCN0000011522) | non-targeting shRNA construct pLKO.1 shSCR (Plasmid #17920) |
| **GPR110 Overexpression** | GPR110 gene: pLV[Exp]-Bsd-EF1A>hADGRF1[NM_153840.4], vectorbuilder, ID:VB900160-9358wfp | empty pLX304 (Plasmid #25890) |
| **BMAL1 luciferase system** | pLV6-Bmal-luc (Addgene plasmid # 68833) |  |
| **Helper plasmid for packaging** | psPAX2; pMD2. G |  |
| **Mitochondrial calcium biosensor** | pMOS028: Mitycam calcium sensor (Addgene plasmid # 163046) |  |
| **Endoplasmic** | pMOS003: GCaMPer calcium sensor (addgene |  |

|  |  |
| --- | --- |
| **reticulum**<br>**calcium**<br>**biosensor** | Plasmid #163047) |
| **Primary**<br>**antibody** | GPR110 Polyclonal Antibody, thermofisher, PA5-100954 |
| **Primary**<br>**antibody** | BMAL1 Monoclonal Antibody (OTI3G9), thermofisher, MA5-25133 |
| **Secondary**<br>**antibody** | Anti-mouse IgG, HRP-linked Antibody (New England Biolabs) |
| **Secondary**<br>**antibody** | FITC-conjugated Goat anti-Rabbit IgG (H+L) (AS011) (abclonal). |

### Animals and Experimental Design

The animal model and experimental procedures have been described previously (43). Briefly, sixteen 33-week-old C57BL/6J mice (8 males and 8 females; The Jackson Laboratory, Bar Harbor, ME, USA) were individually housed with running wheels and maintained under either a standard 12:12 h light-dark cycle (LD) or constant light exposure (LL) (n = 4 per sex per group). Animals had ad libitum access to food and water throughout the experiment.

To induce circadian stress, mice were exposed to LL conditions for 15 days, whereas control animals remained under LD conditions. Circadian timing was determined using wheel-running activity monitoring. Following experimental exposure, tissues were collected at CT/ZT11 and snap-frozen for subsequent analyses. The present study utilized liver samples obtained from this previously described cohort for additional molecular and biochemical analyses.

### Cell culture

To develop a cellular model of circadian rhythm disruption, we employed the CRISPR-Cas9 gene-editing system obtained from Santa Cruz Biotechnology. Specifically, we used a double-nickase plasmid, sc-400808-NIC-2: BMAL1 Double Nickase Plasmid (h2) R: gtgtggattgcaaccgcaaa, F: taccaagagagctggaaagc. which introduces paired cuts to enhance editing specificity. Cells were transfected with the CRISPR construct using Lipofectamine 3000 (Catalog number L3000001), alongside a control double-nickase plasmid as a negative control Control Double Nickase Plasmid (sc-437281). Following transfection, cells were subjected to puromycin selection (2 µg/mL). From this pool, individual clonal populations were isolated. Genomic DNA was extracted from each colony and Sanger sequencing (SANGER sequencing platform; CHU de Québec-Université Laval Research Center) confirmed targeted gene editing at the expected locus.

### Western Blot

Protein expression was analyzed by Western blotting. Cells were washed with cold phosphate-buffered saline (PBS) and lysed in RIPA buffer containing protease and phosphatase inhibitor cocktails. Lysates were incubated on ice for 30 min and clarified by centrifugation at 12,000 × g for 15 min at 4 °C. Protein concentrations were determined using the bicinchoninic acid (BCA) assay according to the manufacturer’s instructions. Equal amounts of protein (30 µg) were mixed with Laemmli sample buffer, denatured at 95 °C for 5 min, and separated by SDS polyacrylamide gel electrophoresis (SDS-PAGE). Proteins were then transferred onto polyvinylidene difluoride (PVDF) membranes using a Semi-dry transfer system. Membranes were blocked with 5% non-fat milk in Tris-buffered saline containing 0.1% Tween-20 (TBST) for 1 h at room temperature and incubated overnight at 4 °C with the appropriate primary antibodies diluted in blocking buffer. After washing with TBST, membranes were incubated with horseradish peroxidase (HRP)-conjugated secondary antibodies for 1 hours at room temperature. Protein bands were visualized using an enhanced chemiluminescence (ECL) detection system and imaged using a chemiluminescence imaging platform. Band intensities were quantified using ImageJ software, and target protein levels were normalized to the corresponding loading control (β*-actin)*. Subsequently, Among the screened clones, two knockout clones were identified, exhibiting complete loss of protein expression. These clones were selected for further functional analyses.

### Immunocytochemistry (ICC)

Cells were seeded onto culture plates (µ-Slide 8 Well high, ibidi: Cat.No:80806) and allowed to attach overnight. The following day, cells were fixed using 4% paraformaldehyde containing Tween-20 (% 0.1) for 20 min at room temperature. After fixation, cells were washed three times with phosphate-buffered saline (PBS) and incubated overnight at 4°C with a primary antibody against GPR110. The following day, cells were incubated with the corresponding fluorescent secondary antibody together with Hoechst nuclear stain for visualization of nuclei. Cells were subsequently washed five times with PBS (15 min each). Fluorescent images were acquired using a confocal microscope (LSM800 microscope, Zeiss Axio Observer Z1 motorized inverted microscope) with a 63× objective lens. For image analysis, fluorescence intensity was quantified using ImageJ software. Individual cells were selected as regions of interest (ROIs), and integrated density values were measured on a per-cell basis. Quantified fluorescence values were exported for statistical analysis. Statistical comparisons between groups were performed using Student’s t-test or analysis of variance (ANOVA), as appropriate.

### Lentiviral Production and Generation of Stable Cell Lines

Lentiviral particles were generated by transient transfection of HEK293T cells with the lentiviral transfer plasmid together with the packaging plasmids psPAX2 and pMD2. G. Transfections were performed using a Lipofectamine 3000 transfection (Thermofisher: Cat: L3000001) reagent according to the manufacturer’s instructions. Viral supernatants were collected 48-72 h post-transfection, clarified by centrifugation, and used to transduce target cell lines.

Following cell line were packaged and generated with responsible construct including pLV6-Bmal-luc, short hairpin RNA, mitochondrial and endoplasmic reticulum Ca²□ Biosensor, GPR overexpression, and appropriate controls were generated at a same time and after 5 passage were stocked in liquid nitrogen (-180 degree) for downstream experiments (each stock was used once per experiment and did not experience any passage after thaw).

For Ca²□ imaging experiments, lentiviral vectors encoding genetically encoded Ca²□ biosensors were used to generate stable reporter cell lines. Following transduction, biosensor expression was confirmed by fluorescence microscopy, and 100 % transduced cell populations were used for downstream analyses. One biosensor was targeted to mitochondria (pMOS028: Mitycam Ca²□ sensor (Plasmid #163046)) to monitor mitochondrial Ca²□ flux (94), while a second biosensor was targeted to the endoplasmic reticulum (ER) (addgene: pMOS003: GCaMPer Ca²□ sensor (Plasmid #163047)) to assess ER Ca²□ dynamics.

To investigate the role of GPR110, stable cell lines with either GPR110 knockdown or ectopic GPR110 expression were generated using lentiviral transduction. For knockdown experiments, cells were transduced with pool of lentiviral vectors expressing (Brown institute, USA) short hairpin RNA (GPR110 (pLKO.1) shRNA:TRCN0000011518, TRCN0000011519, TRCN0000011520, TRCN0000011521, TRCN0000011522) targeting GPR110, while control cells received a non-targeting shRNA construct pLKO.1 shSCR (Plasmid #17920) Scrambled hairpin (negative control, shSCR). For over expression cell line, cells were transduced with lentiviral vectors encoding the full-length GPR110 gene. Following transduction, the expression levels were validated by quantitative PCR and ICC prior to functional experiments.

### Validation of Circadian Rhythm Disruption Using a BMAL1-Luciferase Reporter Assay

To investigate the impact of gene knockout on circadian rhythm regulation, we employed a BMAL1-luciferase reporter construct, provided from Addgene (pLV6-Bmal-luc (Plasmid #68833)). This reporter system consists of the BMAL1 promoter driving luciferase expression, enabling real-time assessment of circadian oscillations. For circadian synchronization, cells were treated with 50% horse serum for 2 hours. Next, luciferase activity was performed and measured at 0, 6, 12, and 24 hours post-synchronization to track rhythmic fluctuations in BMAL1 expression. Because REV-ERB and ROR act as opposing regulators of BMAL1 transcription, their coordinated activity contributes to oscillatory promoter dynamics over time. Specifically, luciferase activity exhibited temporal fluctuations characterized by an initial decrease followed by a later increase, reflecting rhythmic regulation of BMAL1 promoter activity.

### Quantitative Real-Time PCR (TaqMan Gene Expression Assay)

Total RNA was extracted from cells/tissues using TRIzol reagent (Invitrogen, Thermo Fisher Scientific) according to the manufacturer’s instructions and eluted in nuclease-free water. RNA concentration and purity were determined using a BioTek Take3 plate by measuring absorbance at 260 and 280 nm (A260/A280 ratio), and RNA integrity was verified by agarose gel electrophoresis. Complementary DNA (cDNA) was synthesized from 1 µg of total RNA using a High-Capacity cDNA Reverse Transcription Kit (Thermo Fisher Scientific; Cat. No. 4368814) in a total reaction volume of 20 µL, following the manufacturer’s protocol.

Gene expression analysis was performed using TaqMan Gene Expression Assays (Applied Biosystems) on a real-time PCR system (Applied Biosystems). Each reaction was carried out in a 20 µL final volume containing TaqMan Universal PCR Master Mix, gene-specific TaqMan probe, and cDNA template corresponding to ∼500 ng of input RNA. All reactions were performed in technical duplicate.

The thermal cycling conditions consisted of an initial enzyme activation step at 95 °C for 10 min, followed by 40 cycles of denaturation at 95 °C for 15 s and annealing/extension at 60 °C for 1 min. Relative gene expression levels were calculated using the 2^−ΔΔCt method. Expression values were normalized to the endogenous reference genes GAPDH and ACTB, which were selected based on their stable expression across experimental conditions. Results are presented as fold change relative to the control group.

### Gene expression

Total RNA was extracted from cells using QIAzol reagent according to the manufacturer’s instructions. RNA concentration and purity were determined spectrophotometrically, and complementary DNA (cDNA) was synthesized from 500 ng of RNA using the High-Capacity Reverse Transcription Kit (Thermo Fisher Scientific). Quantitative real-time PCR (qPCR) was performed using Abclonal 2× Universal SYBR Green qPCR Master Mix on a real-time PCR system. Amplification was carried out under the following cycling conditions: an initial denaturation step at 95°C for 3 min, followed by 40 cycles of 95°C for 10 s and 60°C for 30 s. Gene expression levels were normalized to the selected housekeeping gene and calculated using the comparative Ct (ΔΔCt) method. qPCR was applied on provided primer sets (Supplementary Table 1).

### Flow Cytometry

HepG2 cells were seeded at a density of 20,000 cells per well and allowed to adhere overnight under standard culture conditions (37 °C, 5% CO□). Cells stably expressing mitochondrial- or endoplasmic reticulum (ER)-targeted Ca²□ biosensors were used for analysis. Following treatment, cells were washed with phosphate-buffered saline (PBS), detached using trypsin-EDTA, and resuspended neutralized in FBS and washed and kept in PBS for analysis. Cell suspensions were analyzed using a BD Biosciences flow cytometer. Fluorescence signals corresponding to Ca²□ levels were detected in the Alexa Fluor 488-A channel. A total of 20,000 events were recorded per sample. Data were analyzed using FlowJo, and gating was performed to exclude debris and doublets.

To assess the effect of docosahexaenoyl ethanolamide (DHEA), cells were treated with 100 nM DHEA. Control cells received vehicle (DMSO, 0.01% final concentration, 1:10,000 dilution). For comparisons between control and BMAL1 knockout (KO) cell lines, Ca²□ levels were measured under basal (untreated) conditions.

### Plate Reader Assay

For fluorescence-based quantification of intracellular Ca²□ levels, HepG2 cells expressing organelle-targeted Ca²□ biosensors were seeded at a density of 12,000 cells per well in 48 well plate and allowed to adhere overnight. Cells were treated with 100 nM DHEA for 2 hours. Following treatment, fluorescence intensity was measured using a microplate reader equipped with appropriate excitation/emission settings (Ex: 488 nm, Em: 515 nm). Background fluorescence was subtracted, and values were normalized by MTT assay and to control conditions.

### AdipoRed assay

To investigate the effect of DHEA on lipid accumulation and to compare lipid content between BMAL1 knockout (BMAL1-KO) and control HepG2 cells, an AdipoRed assay was performed. HepG2 cells were treated with oleic acid (OA, 200 µM) in the presence or absence of DHEA. Following overnight incubation, intracellular neutral lipid accumulation was quantified using the AdipoRed assay. Fluorescence values were normalized to MTT measurements to account for differences in cell viability and metabolic activity.

### Seahorse XF Mito Stress Test

Mitochondrial function was assessed using the Seahorse XF Cell Mito Stress Test assay (Agilent Technologies) according to the manufacturer’s instructions. HepG2-derived cell lines, including wild-type control cells, BMAL1 knockout (KO) cells, shRNA-mediated knockdown cells, and GPR110 overexpressing cells, were used for analysis.

Cells were seeded in Seahorse XF cell culture microplates at an optimized density (typically 20,000-30,000 cells per well) and allowed to adhere overnight under standard culture conditions (37 °C, 5% CO). Prior to the assay, cells were treated with docosahexaenoyl ethanolamide (DHEA, 100 nM) for varying durations. Control wells received vehicle (DMSO, 0.01% final concentration). On the day of the assay, culture medium was replaced with Seahorse XF assay medium (non-buffered DMEM supplemented with glucose, pyruvate, and glutamine, pH 7.4), and cells were equilibrated in a non-CO incubator at 37 °C for 45-60 minutes prior to measurement. Oxygen consumption rate (OCR) was measured under basal conditions and following sequential injections of mitochondrial inhibitors: oligomycin (ATP synthase inhibitor), carbonyl cyanide-p-trifluoromethoxyphenylhydrazone (FCCP; mitochondrial uncoupler), and a mixture of rotenone and antimycin A (complex I and III inhibitors), using the Seahorse XF Analyzer (Agilent Technologies). Mitochondrial respiration parameters, including basal respiration, ATP-linked respiration, maximal respiration, proton leak, and spare respiratory capacity, were calculated according to standard Seahorse protocols. OCR values were normalized to protein content (BCA assay) and cell number.

### Pyruvate metabolism

To investigate the effect of DHEA on cellular pyruvate metabolism, intracellular pyruvate concentrations were measured in different HepG2 cell models, including control cells, BMAL1 knockout (BMAL1-KO) cells, GPR110 knockdown (sh-GPR110) cells, and GPR110 overexpressing (OE-GPR110) cells. Cells were treated with DHEA for 2 hours, after which intracellular pyruvate levels were quantified using a pyruvate assay according to the manufacturer’s instructions (Pyruvate-Glo™ Assay, Catalog number: J4051). The objective was to determine whether DHEA modulates pyruvate levels and to assess whether these effects are mediated through GPR110 signaling and altered circadian regulation. The values were normalized to BCA.

### LC-MS and Lipid Extraction Protocol

Lipid extraction from tissue and plasma samples was performed as previously described using a modified Bligh and Dyer protocol (95). Briefly, tissue samples were homogenized in Tris-HCl buffer (50 mM, pH 7), mixed with methanol containing deuterated internal standards and chloroform, and subjected to liquid-liquid extraction. The organic phases were collected, pooled, evaporated to dryness, and reconstituted in LC-MS mobile phase prior to analysis. Plasma samples were extracted as previously described (96). Lipid mediators were quantified using a Shimadzu 8050 triple quadrupole mass spectrometer as previously reported (97). Retention times were validated using authentic standards injected with each analytical batch. For monoacylglycerols (MAGs), signals from sn-1(3) and sn-2 isomers were combined due to acyl migration during sample handling. Only peaks with a signal-to-noise ratio ≥ 5 were included for quantification (98).

### Cytokines analyses

Cytokine and chemokine levels were determined via Human Cytokine Proinflammatory Focused 15-Plex Discovery Assay® Array (HDF15) (Calgary, AB).

### Limitation

A limitation of the present study is that BMAL1 deficiency does not exclusively reflect circadian clock disruption. BMAL1 has functions beyond the regulation of circadian rhythmicity, and therefore some phenotypes observed in BMAL1 KO cells may result from loss of BMAL1-dependent pathways independent of clock function. Consequently, our findings should be interpreted as effects associated with BMAL1 deficiency rather than definitive evidence of circadian disruption alone. Another limitation of this study is that some Ca²□ measurements were performed with a relatively small number of biological replicates (n = 3). The flow cytometry-based Ca²□ assay required several cell-processing steps prior to analysis, including cell detachment by trypsinization, centrifugation, washing, and resuspension. These procedures may introduce cellular stress and have the potential to influence intracellular Ca²^+^ homeostasis. Although all experimental groups were processed identically to minimize technical variability, the combination of a limited sample size and extensive sample preparation may have reduced the sensitivity of the assay and should be considered when interpreting the Ca²^+^-related findings.

## Author contribution

PAP contributed to performed experiments, analysis and manuscript writing; TDN contributed to performed experiments, edited manuscript; ED contributed to analysis; DM, analysis and manuscript editing; AV, analysis and manuscript editing; NF, analysis and manuscript editing; VDM contributed to analysis and manuscript editing; EMM contributed to experimental design and manuscript editing; and CS contributed to experiment conceptualization, experimental design, manuscript editing, and funding.

## Acknowledgments

This work was supported by the Sentinel North Partner Research Chair on the Gut Microbiome-Endocannabinoid System as an Integrator of Extreme Environmental Influences on Bioenergetics and by the National Institutes of Health (NIH) Grant R15GM134528. The authors thank the members of the laboratory for helpful discussions and technical assistance. We are also grateful to the staff of the institutional animal facility and analytical core platforms for their support with animal care, microbiome sequencing, and lipidomic analyses.

## Abbreviations

ECS: endocannabinoid system
eCBome: endocannabinoidome
NAE: *N*-acylethanolamine
2-MAG: 2-monoacylglycerol
2-AG: 2-arachidonoylglycerol
AEA: anandamide
PEA: *N*-palmitoyl-ethanolamine
OEA: *N*-oleoyl-ethanolamine
DHEA: *N*-docosahexaenoyl-ethanolamine
DHA: docosahexaenoic acid
EPA: eicosapentaenoic acid
SDA: stearidonic acid
LC-MS/MS: liquid chromatography-tandem mass spectrometry.

