## Supplementary figures and images for "Circadian disruption alters hepatic calcium hemostasis, endocannabinoidome and mitochondria through *N*-docosahexaenoyl ethanolamide-GPR110 signaling"

### supplementary figures

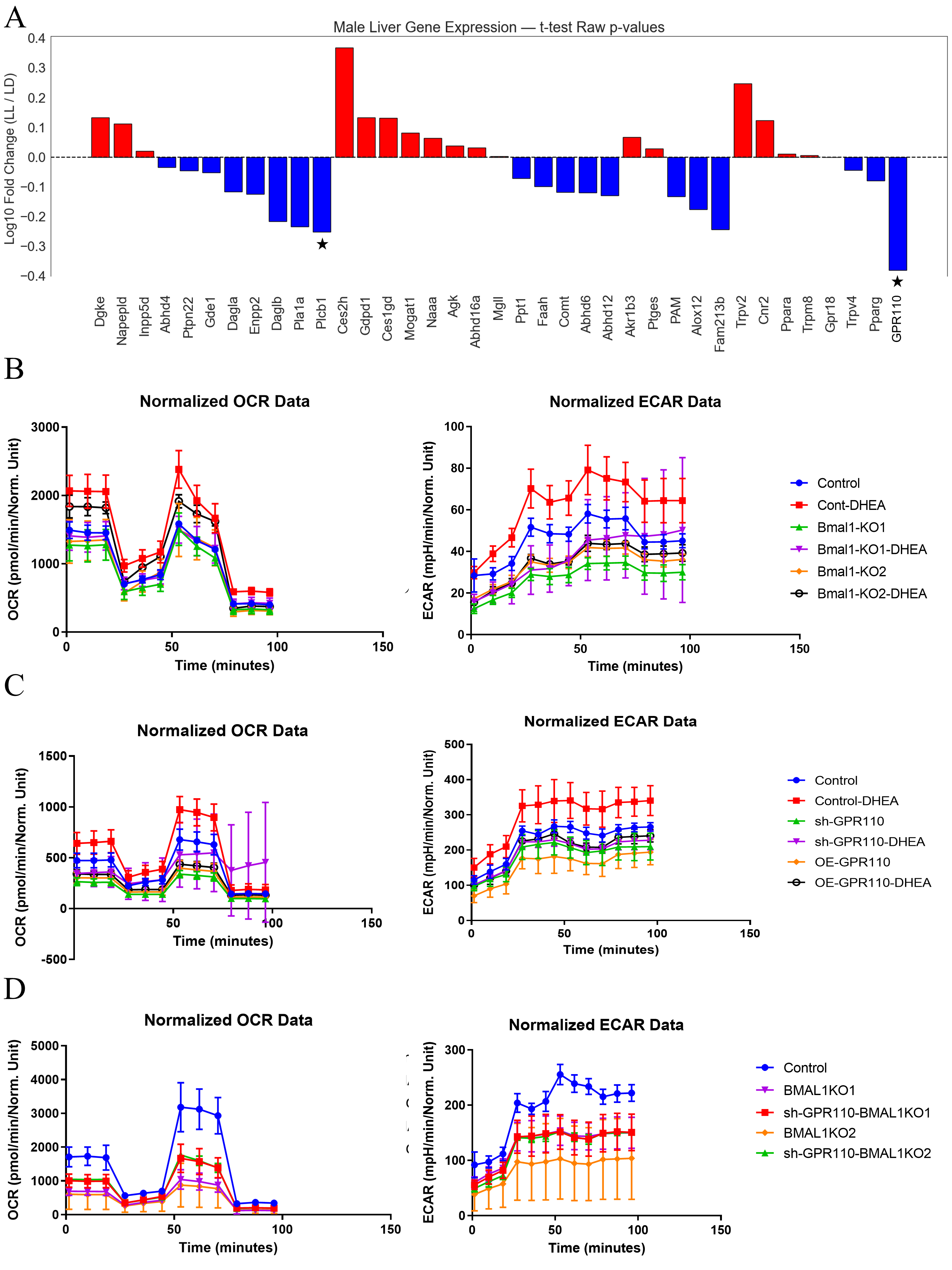
